# Mathematical modeling of tumor-tumor distant interactions supports a systemic control of tumor growth

**DOI:** 10.1101/168823

**Authors:** Sebastien Benzekry, Clare Lamont, Dominique Barbolosi, Lynn Hlatky, Philip Hahnfeldt

## Abstract

Interactions between different tumors within the same organism have major clinical implications, especially in the context of surgery and metastatic disease. Three main explanatory theories (competition, angiogenesis inhibition and proliferation inhibition) have been proposed but precise determinants of the phenomenon remain poorly understood. Here we formalized these theories into mathematical models and performed biological experiments to test them with empirical data. In syngeneic mice bearing two simultaneously implanted tumors, growth of only one of the tumors was significantly suppressed (61% size reduction at day 15, p<0.05). The competition model had to be rejected while the angiogenesis inhibition and proliferation inhibition models were able to describe the data. Additional models including a theory based on distant cytotoxic log-kill effects were unable to fit the data. The proliferation inhibition model was identifiable and minimal (4 parameters), and its descriptive power was validated against the data, including consistency in predictions of single tumor growth when no secondary tumor was present. This theory may also shed new light on single cancer growth insofar as it offers a biologically translatable picture of how local and global action may combine to control local tumor growth, and in particular, the role of tumor-tumor inhibition. This model offers a depiction of concomitant resistance that provides an improved theoretical basis for tumor growth control and may also find utility in therapeutic planning to avoid post-surgery metastatic acceleration.

## Major findings

In mice bearing two tumors implanted simultaneously, tumor growth was suppressed in one of the two tumors. Three theories of this phenomenon were advanced and assessed against the data. As formalized, a model of competition for nutrients was not able to explain the growth behavior as well as indirect, angiogenesis-regulated inhibition or a third model based on direct systemic inhibition. This last model offers a depiction of concomitant resistance that provides an improved theoretical basis for tumor growth control and may also find utility in therapeutic planning to avoid postsurgery metastatic acceleration.

## Quick guide to equations and assumptions

Volumes of the two tumors at time *t* were denoted *V*_1_(*t*) and *V*_2_(*t*) and differential equations were derived for the rate of change of these quantities.

For biological relevance, we required that the model(s) comply with the following conditions: 1) in the absence of a second tumor, the model had to be able to fit single-tumor growth curves, 2) the shape of the inhibition effect had to be identical from tumor 1 on tumor 2 as from tumor 2 to tumor1 (structural symmetry) and 3) the parameters had to be identical for the two tumors (parametrical symmetry). The source of the observed difference between the two growing tumors was assumed to result from an initial (small) discrepancy in the number of cells that take during the tumors grafts, respectively denoted *V*_0,1_ and *V*_0,2_. After investigation of the sensitivity of several models to these quantities and their ratio, we considered more relevant for robustness of the results to fix their ratio for all the animals. It was set to a twenty-five percent higher cell loss in the inhibited tumor, as compared to the non-inhibited one. We thus fixed *V*_0,1_ = 1 mm^3^ (≃ 10^6^ cells (1)) and *V*_0,2_ = 0.75 mm^3^.

Our main model is based on experimental evidence from (2,3) demonstrating that a tumor produces inhibitory factors (IF), such as meta- and ortho-tyrosines, that induce a cell cycle arrest (Figure 1).

**Figure 1:**
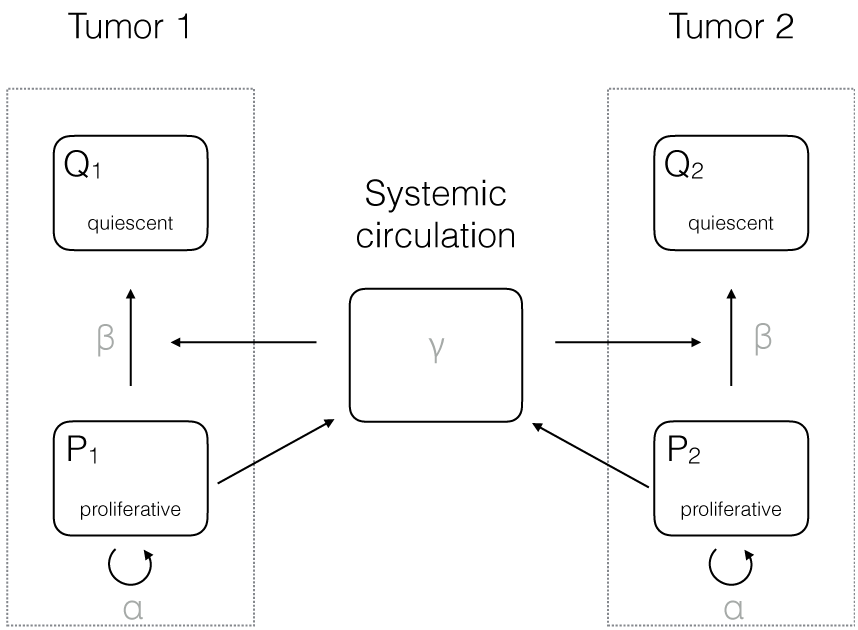
Scheme of the proliferation-inhibition model.

The model assumptions are:

- Proportionality between volume and number of cells, using the well-established conversion rule 1 mm^3^ ≃ 10^6^ cells.
- Each tumor volume is divided into two compartments: proliferative cells (*P_i_* with *i* the tumor index) and non-proliferative tissue (*Q_i_*).
- Proliferative cells have a constant length of the cell cycle *τ*. The proliferation rate 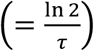 is denoted *α* (day^-1^).
- Proliferative cells release IFs with a rate proportional to their number. The proportionality coefficient is denoted *β*_0_ and is expressed in mol × mm^-3^ × day^-1^. There is a local elimination of IFs with rate *k_loc_* (day^-1^) and the concentration is assumed to be at steady state.
- A fraction *ϕ* of these factors is released into the systemic circulation.
- In the blood, IFs are subject to a first order elimination process with rate *k* (day^-1^). We assume that the time scale of the blood distribution is faster than the tumor growth and thus consider the blood concentration at steady-state. Assuming that a fraction *ψ* reaches the distant site, the concentration of IFs at the distant site is therefore 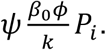 At the local site it is 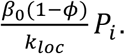
- At each local site, the IFs induce a proliferation arrest, making cells go from a proliferative state to a quiescent state. A given amount of these molecules provokes cell cycle arrest of a constant *number* of proliferative cells (in contrast to a constant *fraction* in the log-kill law usually employed for cytotoxic agents (4)), with rate *β*_1_ (mol^-1^ × mm^3^ × day^-1^). Denoting 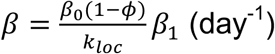 and 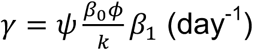 the number of cells going from a proliferative state to a quiescent state within the tumor *i* is thus *β P_i_* + *γ*(*P_i_* + *P_j_*). Note that this includes both local inhibition and global inhibition, which accounts for factors released by the other tumor (tumor *j*) but also tumor *i* itself.

The model then writes:

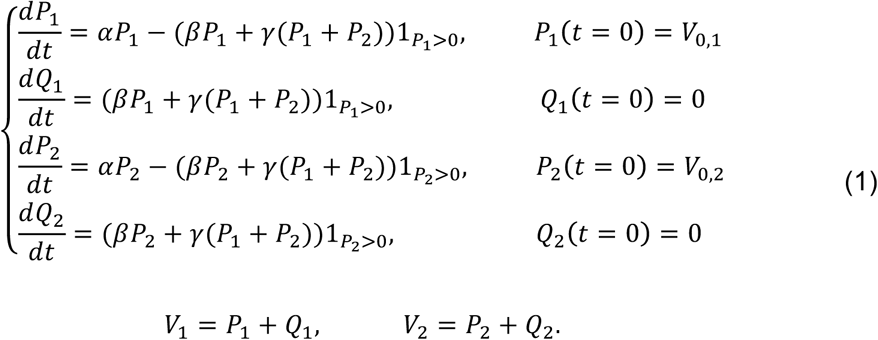

The Heaviside functions 1_*P*_1_>0_ and 1_*P*_2_>0_ (equal to one if *P_i_* > 0 and zero elsewhere) stand for the fact that when factors are present but no proliferative cells exist, no cells go to quiescence. In particular, they ensure that the solutions (understood in the weak sense due to the discontinuous nature of the Heaviside function) remain positive.

## Introduction

Concomitant tumor resistance (CR) is a phenomenon by which the presence of a tumor negatively influences the appearance and growth of another implant (see (5) for a review). It has been reported in numerous experimental studies spanning over century, and implementing a large variety of animal models (6–16). These studies investigated either the concomitant or subsequent implantation of a second graft after a primary injection (7,9,10,12,13,16), or the inhibition of secondary tumors arising from the primary (metastases) (8,17–20). They consistently evidenced a systemic growth suppression effect, demonstrated by the occurrence of post-removal growth acceleration (9,10,14,17,18,20–23). In the clinic, suppression of the growth of metastases by the presence of the primary tumor has yet to be appreciated in general therapeutic planning, although it has been reported in patients (21,24–26). Despite the abundance of reports of this phenomenon, the precise determinants of CR remain poorly understood and only qualitative theories have been advanced.

CR is of direct clinical relevance insofar as it implies that removal of a primary tumor, with the resultant release of its inhibitory pressure on occult secondary sites, could be followed by post-surgery metastatic acceleration (PSMA). PSMA has been demonstrated to occur in numerous animal experiments (9,10,17,20,23), as well as in clinical case reports (24,25,27). Further support for the occurrence of PSMA in a notable fraction of patients was also provided by the observation of two peaks in the hazard relapse rate of a large cohort of breast cancer patients (28,27,26,29).

Several hypotheses have historically been proposed for the explanation of the underlying mechanism of CR. The first was due to P. Ehrlich and consisted in *athrepsia*, i.e. that two (or more) tumors in the same organism would compete for nutrients and that the growth of one tumor would leave less nutrients available for the other (11,30). However, this was challenged by the observations that CR was decreased when the number of inoculated cells was increased (13). Another popular theory, first introduced by Bashford in 1908, was based on immunologic mechanisms and stipulated that the presence of a first tumor would activate an immune response preventing the second graft to take or grow (7,11). However, several studies demonstrated the occurrence of CR in tumor models with no or weak immunogenicity, or in immune-deprived mice, thus challenging this explanation (8,11–13). This implies that, although immunologic factors might contribute to CR, they cannot explain it entirely.

In the 1990’s, a team led by J. Folkman discovered endogenous inhibitors of angiogenesis (such as endostatin or angiostatin) by demonstrating that injection of these factors could substitute for the suppressive effect on lung metastases exerted by the primary tumor (18,31). This led the investigators to link CR to distant systemic inhibition of angiogenesis. Their theory is based on the fact that locally produced angiogenesis inhibitors would spread systemically using the vascular system, reach secondary lesions and impair angiogenesis at the distant sites (15,16,21,32), eventually overcoming the influence of angiogenesis stimulators also produced by the tumors. This idea is supported by the fact that endogenous angiogenesis inhibitors have long half-lives compared to stimulators, which allows them to diffuse, reach the vascular system and accumulate (18,33).

The idea of circulating inhibiting factors due to the presence of a primary tumor had been proposed and confirmed in earlier studies (5,8,11), but their precise mode of action has remained elusive. Distinct from the angiogenesis inhibitors previously mentioned, another research group identified other blood-borne factors with direct anti-proliferative action, namely meta- and ortho-tyrosine, that would reduce proliferation by driving tumor cells into a G0-phase state of dormancy or induce an S-phase arrest (2,3).

So far, arguments and theories about CR have remained qualitative in nature. In the present study, comparing alternative mathematical formalisms, we demonstrate that a simplified model with well-motivated parameters that addresses concomitant resistance specifically was able to capture features of coupled tumor growth and may shed more light on the understudied subject of systemic controls in cancer, a potentially critical step toward eventually understanding metastatic control.

## Materials and Methods

### Mice experiments

#### Cell culture

Murine Lewis lung carcinoma (LLC) cells, originally derived from a spontaneous tumor in a C57BL/6 mouse (34), were obtained from American Type Culture Collection (Manassas, VA) in the period 2005-2006. The LLC cells were cultured under standard conditions (34) in high glucose DMEM (Gibco Invitrogen Cell Culture, Carlsbad, CA) with 10% FBS (Gibco Invitrogen Cell Culture) and 5% CO_2_. They were expanded through 4 passages (LLC p4) and then aliquoted and stored in liquid nitrogen. At that time, they underwent Molecular Diagnostics Infectious Disease PCR testing (Mouse Essential Panel) at Charles River Laboratories. An aliquot of the above LLC p4 cell line was thawed and tested for Mycoplasma at Bionique Laboratories for Mycoplasma testing. They were found to be Negative for Mycoplasma. The cells have not been characterized. An aliquot of LLC p4 was used in these experiments. LLC p4 cells were thawed and passaged through an additional 2 cycles prior to injection into the mice.

#### Tumor Injections

C57BL/6 male mice were used with an average lifespan of 878 days (35). At time of injection mice were 6 – 8 weeks old (Jackson Laboratory, Bar Harbor, Maine). Subcutaneous injections of 10^6^ LLC cells in 0.2 ml phosphate-buffered saline (PBS) were performed on the caudal half of the back for the control group and on the two lateral sides of the caudal half of the back for the group bearing two tumors, in anesthetized mice. Tumor size was measured regularly with calipers to a maximum of 1.5 cm^3^ when mice were sacrificed. Animal tumor model studies were performed in strict accordance with the recommendations in the Guide for Care and Use of Laboratory Animals of the National Institute of Health and according to guidelines of the Institutional Animal Care and Use Committee at Steward Research and Specialty Projects Corp.

### Statistical analysis

Simulations of the mathematical models were performed using Matlab with statistics and optimization toolboxes (The Mathworks Inc., 2015). Fits of the models to the data were executed using an internally developed software that utilizes the function *fminsearch* for weighted least squares minimization. The objective function was computed using weights defined by an error variance model previously established on the same animal model and measurement technique (36). Statistical analyses of the fits (computation of the goodness-of-fit metrics and standard errors of the parameter estimates) were implemented in our software as previously described (36,37).

## Results

We studied the phenomenon of CR by combining experiments and mathematical modeling, informed by pre-existing theories in the literature. For the experiments, two groups were considered. The first group (control) consisted of twenty mice in which single implants were performed. In the second group (double tumors, abbreviated as DT) consisting of ten mice, two grafts with identical load (10^6^ cells) were performed on day 0, at the same locations on opposite flanks of the mice.

### In a mouse bearing two tumors, one has normal kinetics and the growth of the other is suppressed

We first performed a direct (i.e., not model-based) statistical analysis of the data. We compared control tumor growth kinetics in mice bearing single implants with the growth curves of tumors in a double-tumor bearing host (Figure 2). Observations of the kinetics of each tumor in the DT group suggested that in each mouse, one tumor was growing faster than the other, possibly inhibiting the second one (Figure 2A and supplementary Figure 1A-B). This behavior was observed consistently in all the animals of the DT group except in two of them (animals 2 and 9 in Figure 2A), and did not seem to result from the lateral location (left or right) of the tumors (Figure 2A). Intriguingly, the two mice where the phenomenon was not observable were found to have a connecting blood vessel joining the two tumors and were the only ones to exhibit this macroscopic vascular structure. Direct sharing of same vasculature seemed to equilibrate tumor expansions. One possible explanation for the absence of cross growth suppression could be that the production of inhibitors was negligible in these tumor-host systems. This could also explain the formation of the connecting blood vessel due to increased neo-angiogenesis (under the theory of angiogenesis inhibition-driven growth suppression).

**Figure 2:**
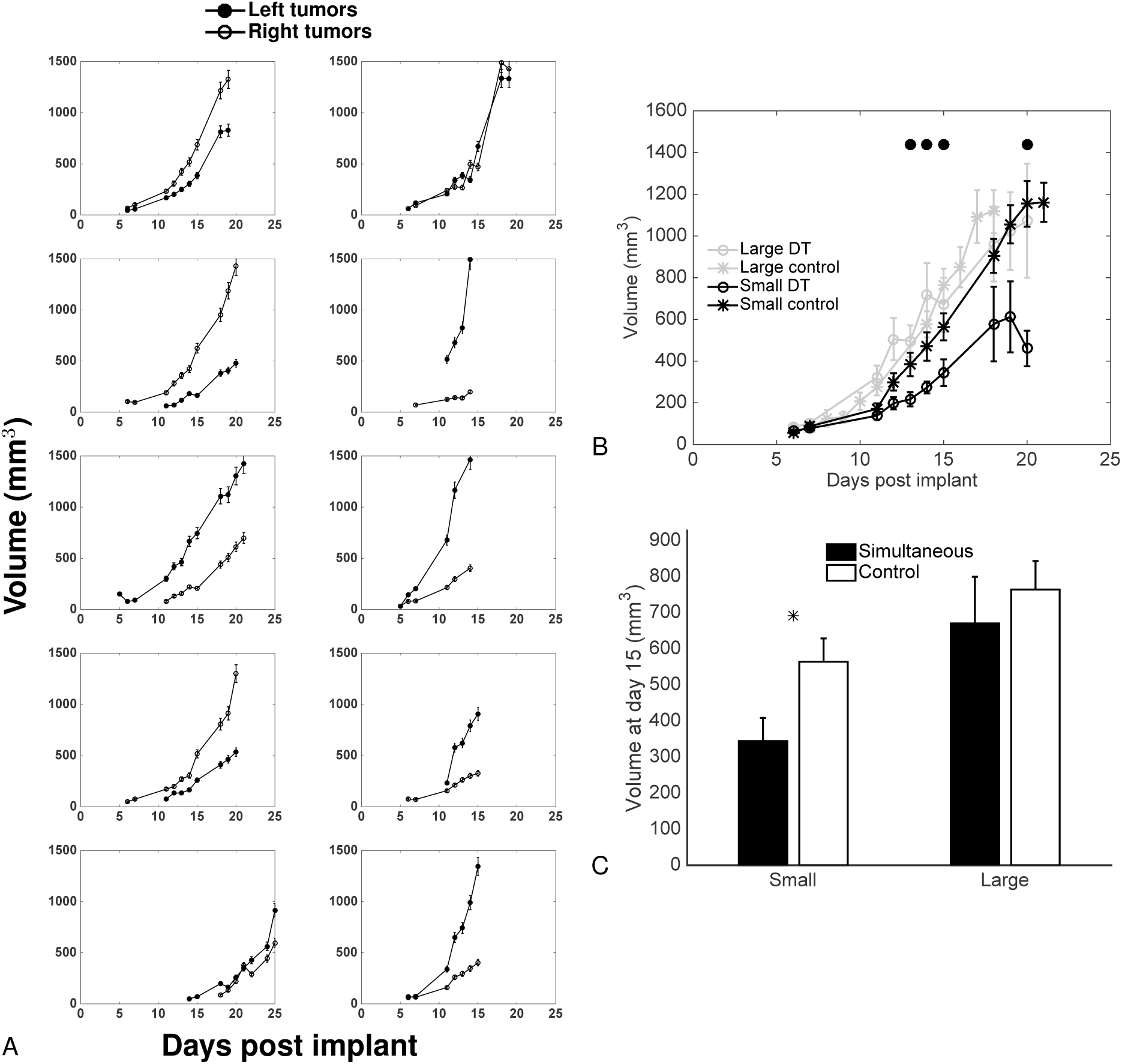
Data of dynamics of simultaneous tumor growth. A. Dynamics of the left and right tumors from mice inoculated with 1×10^6^ LLC cells on the two lateral sides at day 0. B. Comparison of large and small tumors with large and small tumors extracted from artificially paired control tumor growth curves (see text for details). Mean ± standard error. Circles indicate statistically significant differences between the small tumors from the simultaneous group and the small control tumors (Student’s t-test with unequal variance, p<0.05). C. Tumor sizes at day 15. Mean ± standard error. * = p<0.05, Student’s t-test with equal variance.

In order to statistically confirm that this observation was not purely due to intrinsic randomness in experimental conditions (such as the number of initial cells that “take” from the injection) that would by themselves generate different growth curves for the two implants in the same mice, we performed a statistical analysis. It aimed at testing the null hypothesis that both tumors would be identically and independently growing (i.i.g.). We artificially generated 10 couples of i.i.g. tumors by subdividing the control group of 20 animals into two groups of 10, randomly picking tumors from each subgroup and pairing them together. We then picked each small tumor from these pairs (and similarly, each large tumor), by choosing the one with smallest final volume. This yielded two samples of 10 “control small” and “control large” tumors that can be considered as what would have emerged from randomness only in initiation and growth. These two samples could then be statistically compared to the experimental samples of small and large tumors from the DT group. We observed significant differences between the small tumors from the DT group and the “control small” tumors from day 12 until the end (p<0.05, Student’s t-test), with the exception of days 18 and 19 where differences were not significant due to large variability (Figures 2B and 2C). On the other hand, no statistical difference was observed between the large tumors from the DT group and their control counterparts.

These results demonstrate that in a mouse where two tumors are simultaneously growing, the larger tumor is growing at the same speed as would a single tumor, while the other has significantly slower kinetics.

### First insights from modeling: single tumor growth models

To investigate what caused the behavior established in the previous section, we fitted various growth models previously shown able to describe syngeneic LLC single tumor growth with identifiable parameters (36). Models that were considered here were: exponential growth (with free initial volume *V*_0_), i.e. 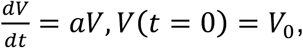 power law, i.e., 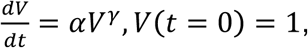 and Gompertz, 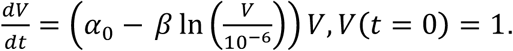 For the two last ones, for identifiability reasons the initial volume was fixed to the number of injected cells (10^6^ cells ≈ 1 mm^3^). The objective was to see if any of these models was able, through estimation of its coefficient, to elucidate the observed differences in the growth of the two tumors in the same animal. In this first context, each tumor was fitted independently, resulting in one parameter set per tumor, and we looked for statistically significant differences in the parameters between the group of small (or large) tumors and the group of small (or large) tumors built from pairings of tumors from the single tumor control group.

As expected from their accurate descriptive power, all the models performed well in terms of goodness of fit and were able to reproduce the tumor growth curves (supplementary Figures 2-4).

Parameter estimates are reported in Figure 3. The only parameter that exhibited significant difference between the simultaneous and control groups was the proliferation rate *α* of the exponential model. This suggests an alteration of proliferative abilities in the altered tumor due to the presence of the other one. On the other hand, the time *t_D_* to reach a given volume *V_D_* (of 100 mm^3^ for instance) was steadily found significantly larger in the small tumors of the two-tumors animals, as compared to the small control tumors (supplementary Figure 5).

**Figure 3:**
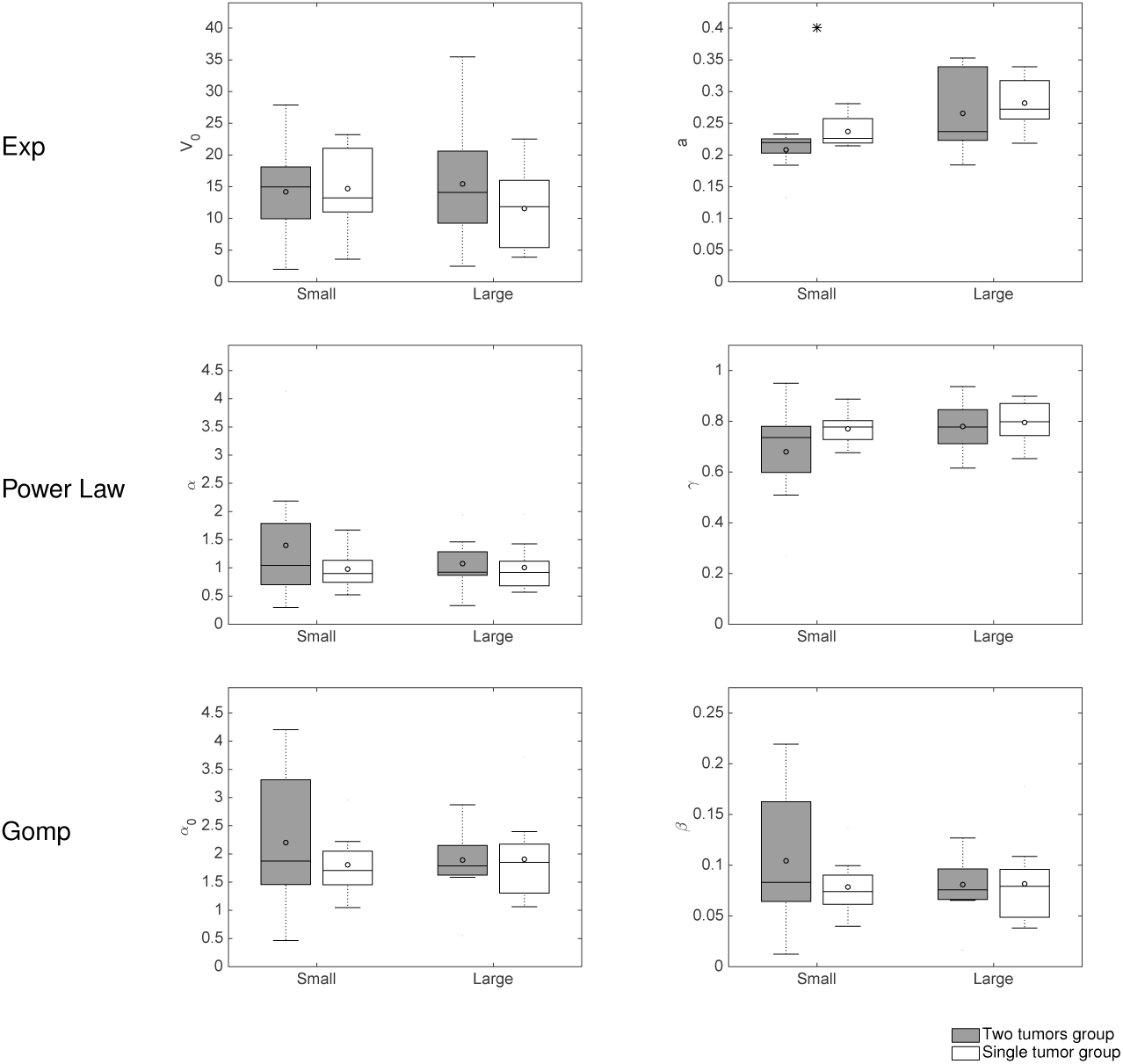
Single-tumor growth models’ analysis: parameter distributions. Models for single tumor growth were independently fitted to the large and small growth curves from the two-tumor bearing animals and from the simulated double-independent tumors from the control group. Shown is a comparison of the distribution of the inferred parameters. Exp = Exponential, Gomp = Gompertz

In summary, single tumor growth models were able to give an appropriate description of the growth curves taken independently and suggest alteration of the proliferative abilities in the small tumors group. However, they don’t offer any insight on the dynamics of interactions between the two implants.

### A dynamical benchmark of models of concomitant resistance

To next study and quantify the possible underlying mechanisms of tumor-tumor interactions leading to the growth kinetics differences that were observed, we investigated models of simultaneous growth of two tumors in the same organism. A virtually infinite number of models can be conceived for description of CR, both in terms of structural shape (equations) and values of the parameters. We report here only on the results from analysis of 5 informative models. Supplementary table 1 details the equations of these models. Interestingly, several models were found unable to fit the data, suggesting rejection of (at least one of) the hypotheses that they rely on. These include a model for the athrepsis theory (competition for nutrients), as shown in supplementary Figure 6. The other models were based on either systemic inhibition of angiogenesis (SIA) – formalized using a model of interaction between tumor growth and vascular support (38) – or induction of quiescence due to a cytostatic seric factor, as experimentally evidenced by Ruggiero et al. (3, 13). We termed the models based on this last theory proliferation inhibition (PI) models. The SIA model was able to give a reasonably accurate description of our data (supplementary Figure 7). For PI models, which have similar structures as equation (1), three hypotheses were investigated (see supplementary Table 1 for the detailed expressions of the models): 1) direct effect (a given quantity of IFs induces a given *number* of cells going to quiescence) (equation (1)), 2) log-kill effect (a given quantity of IFs induces a given *fraction* of cells going to quiescence) and 3) total number of cells (*P_i_* + *Q_i_*) as a source of IFs. Notably, models 2) and 3) were unable to fit the data and had to be rejected (supplementary Figures 8, 9). Residuals analysis supporting these results are shown in the supplementary Figure 10A-D. On the other hand, the model 1) gave a particularly good fit to the data (Figures 4A and B). Table 1 summarizes statistical quantitative metrics of goodness-of-fit the models that allow comparison of their descriptive power, while Figure 4C shows the distribution of the residuals. Table 2 reports the parameter values of all the models estimated from the best fits, together with their inter-animal variability (parameters were individually fitted for each mouse) and standard errors for the estimates.

**Table 1:**
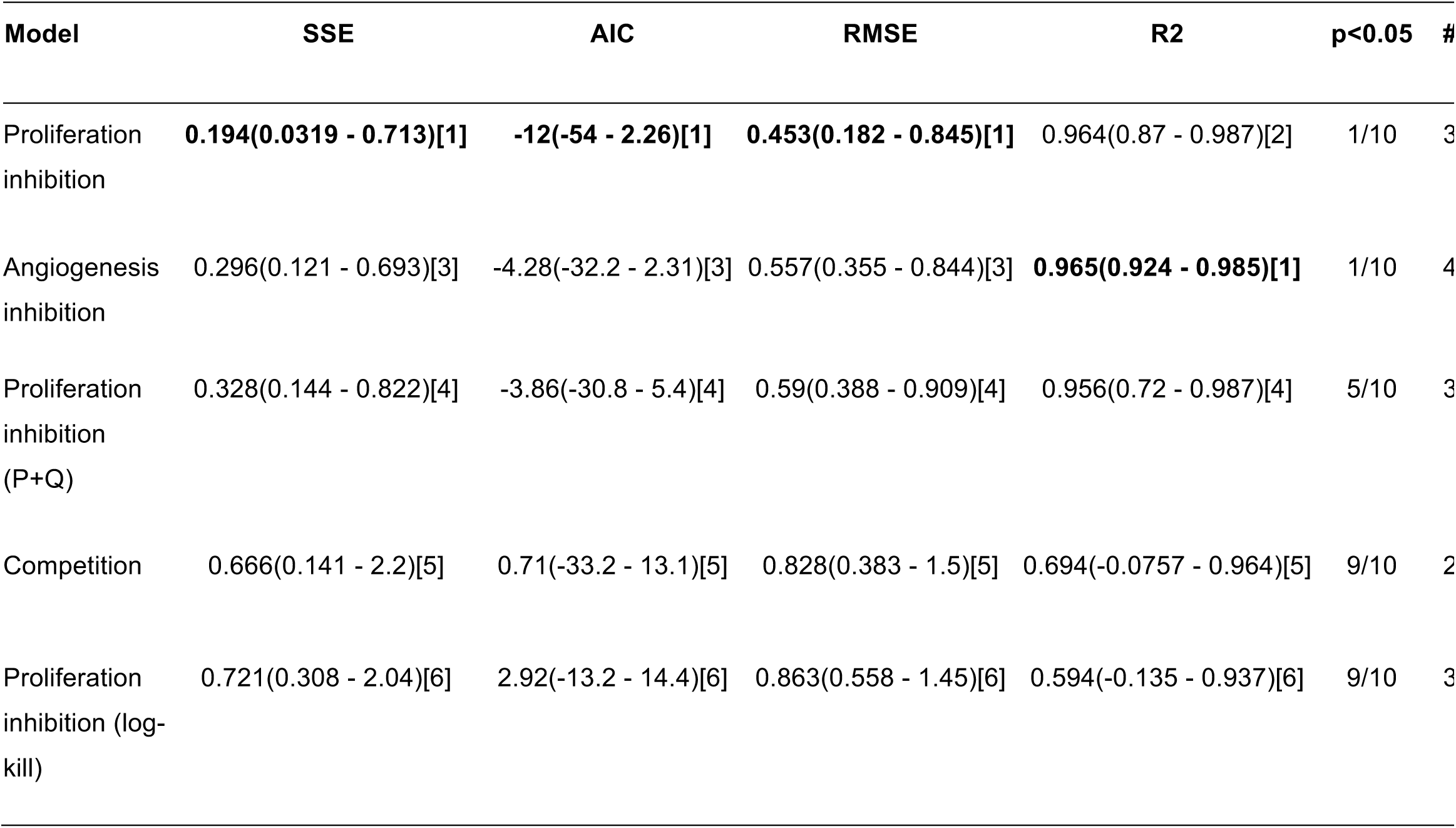
Goodness-of-fit metrics of the two-tumors models. SSE = Sum of Square Errors, AIC = Akaike Information Criterion, BIC = Bayesian Information Criterion, R2 = coefficient of determination. # = number of parameters. The numbers in parentheses indicate the (min - max) range of the values and the numbers inside the brackets are ranks of the models relatively to the criterion (in bold is the model ranking first). The “p<0.05” column contains number of animals for which the null hypothesis of a gaussian distribution of the residuals was rejected for either the large or the small tumor (Kolmogorov-Smirnov goodness-of-fit test).

**Table 2:** Two-tumors models’ parameter values.

| Model | Par. | Unit | Median value (CV) | RSE (%) |
| --- | --- | --- | --- | --- |
| Proliferation<br>inhibition | $\alpha$ | day<sup>-1</sup> | 3.63 (58) | 21.7 |
| | $\beta$ | day<sup>-1</sup> | 3.29 (64.6) | 25.9 |
| | $\gamma$ | day<sup>-1</sup> | 0.0405 (61.2) | 3.94 |
| Angiogenesis<br>inhibition | $a$ | day<sup>-1</sup> | 3.06 (1.06e+03) | 47.4 |
| | $b$ | day<sup>-1</sup> | 0.57 (24.8) | 3.02 |
| | $d$ | mm<sup>-2</sup> · day<sup>-1</sup> | 0.00368 (35.3) | 20.4 |
| | $e$ | day<sup>-1</sup> | 0.119 (43.5) | 1.72 |
| Proliferation<br>inhibition (P+Q) | $\alpha$ | day<sup>-1</sup> | 0.701 (26.9) | 3.61 |
| | $\beta$ | day<sup>-1</sup> | 0.187 (50.1) | 5.31 |
| | $\gamma$ | day<sup>-1</sup> | 0.00754 (253) | 9.45 |
| Competition | $a$ | day<sup>-1</sup> | 0.085 (30.9) | 3.45 |
| | $K$ | mm<sup>3</sup> | 8.36e+03 (2.17e+06) | 15 |
| Proliferation<br>inhibition (log-kill) | $\alpha$ | day<sup>-1</sup> | 0.531 (33.8) | 7.45 |
| | $\beta$ | day<sup>-1</sup> | 7.72e-09 (1.1e+07) | 9.41e+07 |
| | $\gamma$ | day<sup>-1</sup> | 0.00168 (254) | 13.4 |

CV = coefficient of variation. RSE = relative standard error.

**Figure 4:**
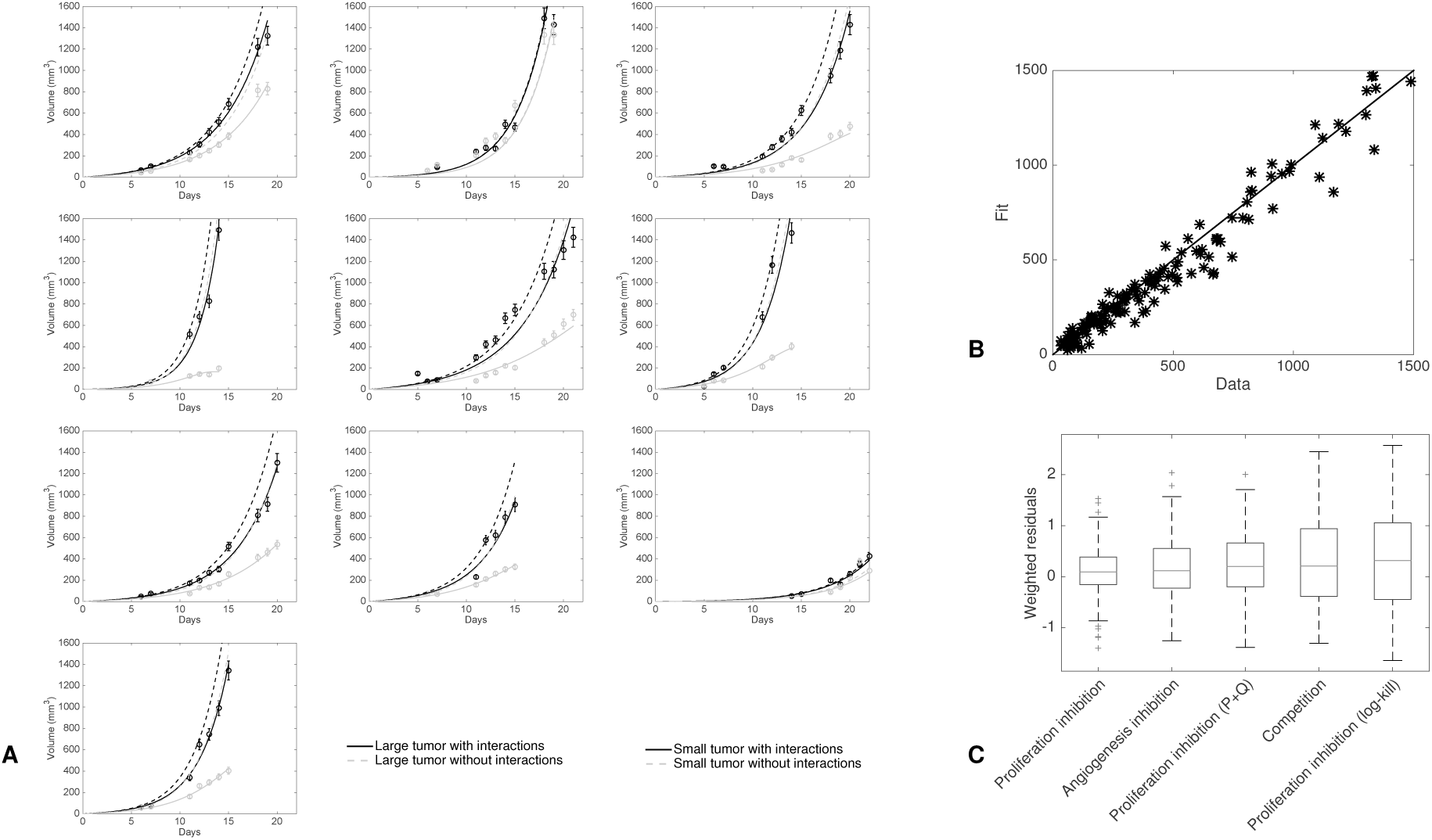
Individual fits of the two-tumors proliferation-inhibition model. A. Fits of all the animals. In each mouse the only difference between the two tumors lies in the number of tumor cells that take, i.e. parameter *V*_*0*,*2*_. Dashed lines are simulations of the model with no interactions between the two tumors (i.e. without terms containing *P*_2_ in the equations on *P*_1_ and *Q*_1_ and conversely for the equations on *P*_2_ and *Q*_2_). B. Fitted values versus data points for all the two-tumors growth curves of A. The solid line is the identity function. C. Distribution of the residuals for the best-fits of the three two-tumors models.

These results demonstrate that a mathematical modeling approach was able, by confrontation of the best-fits of the model, to discriminate among qualitatively equally likely theories of CR and suggest a PI model, having attributes that may explain (or at least describe) this particular phenomenon.

### Validation of a simple and biologically-based mathematical model of CR

#### Double tumor growth

The PI model 1), formalized by equations (1), consists in assuming a direct and mutual growth rate decrease between the two tumors, due to passage to quiescence (Figure 1). Goodness of fit was found excellent (Figure 4), as well as identifiability of the parameters (see standard errors in Table 2). Notably, while being fitted directly on the two tumors growths, the predicted behavior when simulating no interactions was in full agreement with the control growth curves. Indeed, the dashed lines in Figure 4A are close to the growth curves of the large tumors, which were found to be not significantly different from “naturally happening” large tumors. Hence the model was able to learn and identify the unaltered growth part from altered growth curves, highlighting its reliability.

In mouse number two (second plot in the top row of Figure 4A), consistently with the observation of identical growth kinetics between the two tumors, the model identified a value of parameter *γ* not significantly different from zero. On the other hand, the model did identify interactions between the two tumors in the other animals, as emphasized by 95% confidence intervals of parameter *γ* inferred from the parameter estimation process that did not contain 0 ((0.0374, 0.0436) in our estimation). In turn, this translated into substantial differences in the kinetics (see Figure 4A where growth curves are plotted with or without interactions). Of important note, this difference in the kinetics was mostly due to the interaction between the two tumors, rather than the initial difference in cell loss. This is demonstrated in Figure 4A where it can be observed that the growth curves of the small tumors with only a different initial volume (dashed curves) were considerably higher than the curves where interaction was taken into account. Moreover, these curves were both close to the large tumor growth curves, indicating that the difference in *V*_0_ had only a negligible impact on the difference between the two growth curves, the major determinant being the tumor-tumor cross inhibition effect. Critically, the differences for the large tumor curves were much smaller than for the small tumor curves (while the interaction parameter was the same for both tumors), indicating that the model gives a valid quantitative theory of why only one tumor was affected by CR.

Together, our results provide a biologically-based and minimally parameterized mathematical model for tumor growth kinetics interactions in a two-tumors bearing host. The model confirmed a significative non-zero value for the interaction parameter in 9/10 mice, which gave a quantitative measure of the phenomenon. Asymmetry between the two tumors was explained by an initial difference in the take between the two implants.

#### Single tumor growth

In addition to being able to describe double tumor growth and CR, the elementary model that we propose also offers a simple formalism for single tumor volume growth. The model consists in the division of the cancer cells into two subpopulations: proliferative and quiescent cells (which could also account for necrotic tissue still present in the total volume measurement). Indeed, in the absence of a secondary tumor, the model (1) becomes:

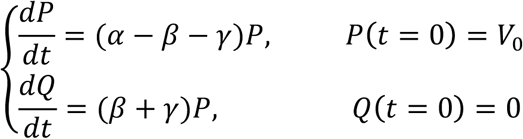

This provides a valid and simple mathematical construct able to describe the growth of single tumors (Figure 5A-B, supplementary Figure 11). It sheds new lights on general tumor growth laws as it demonstrates that classical Gompertzian growth -- which is able to describe accurately Lewis Lung tumor growth curves (36) -- can be reproduced by these equations, with no significant differences (i.e. a difference in Akaike Information Criterion less than 2, see supplementary Table 2). Indeed, it had remained elusive why the Gompertzian curve, which was originally designed not even for growth processes (39), describes tumor growth curves and their consistent relative growth rate decrease with such important accuracy, while being only phenomenological and not biologically grounded. Interestingly, we obtained that a model where growth deceleration was assumed to result (only) from passage to quiescence due to the production of factors by the proliferative tumor cells themselves was able to explain single tumor growth curves as accurately as the Gompertz model, or other models such as the power law (Figure 5A-B). Parameters identifiability of this new model was also very good (supplementary Table 3).

**Figure 5:**
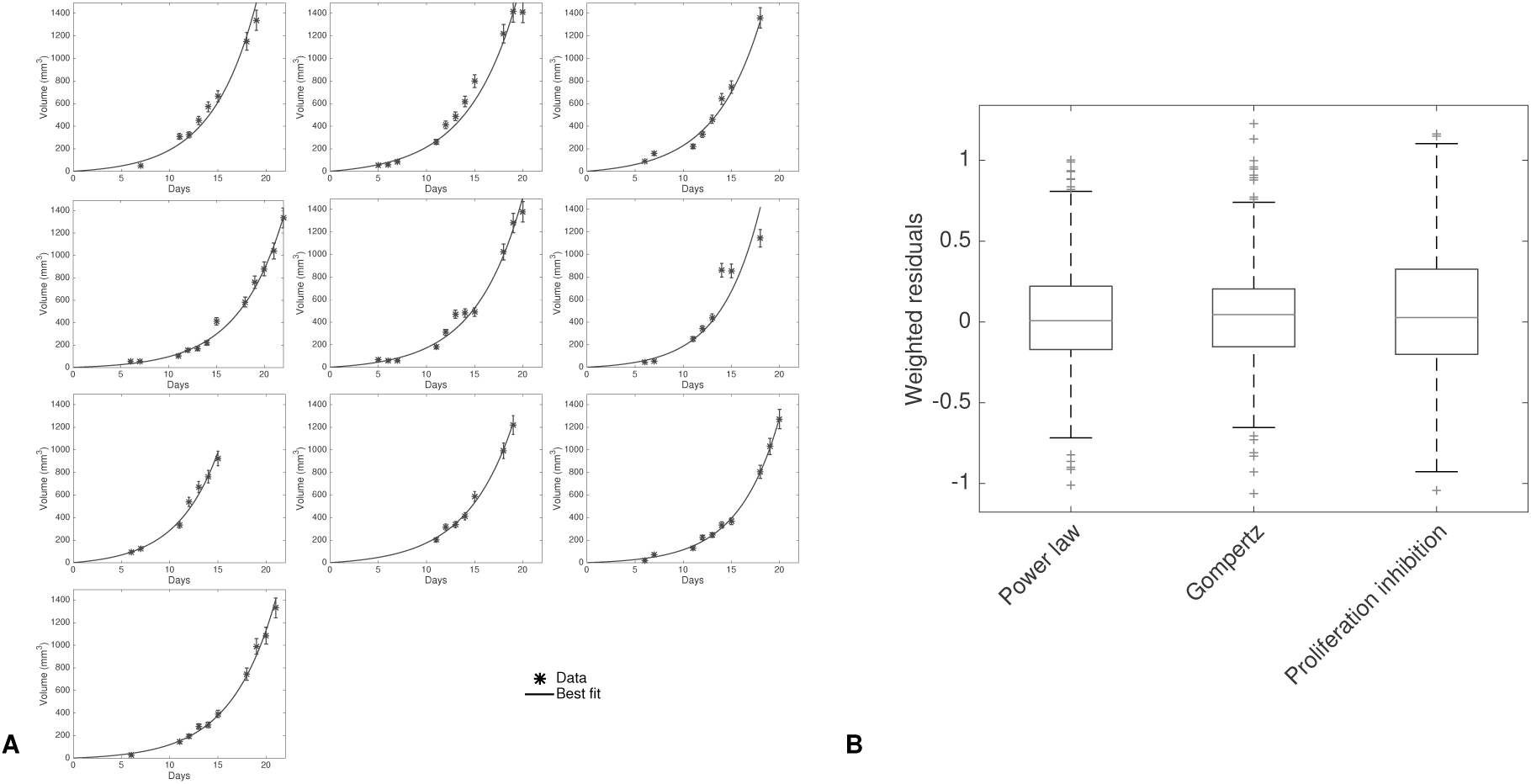
Individual fits of the single tumor proliferation-inhibition model. A. Fits of the single tumor growth model corresponding to model (1) to 10 representative tumor growth curves of the control group. B. Distribution of the residuals for the best-fits of the proliferation-inhibition model and two other classical models of tumor growth: the power law and Gompertz models.

## Discussion

To quantitatively inform on CR, we performed an integrative study that combined experimental and theoretical investigations. At the experimental level, we found that when two identical tumors were implanted in the same organism, distant interactions occurred. Specifically, in one (and only one) of the two tumors, growth was suppressed, while the other remained unaltered. Several theories formalized by mathematical models were assessed against the data by means of appropriate mathematical constructs. Our findings revealed that direct inhibition of proliferation was able to appropriately describe the data. Under this hypothesis, both tumors release systemic factors that mutually induce quiescence in the other tumor, which is in line with experimental results identifying these factors as meta- and ortho-tyrosine (2,3). In our study, the origin of the asymmetry between the two tumors was hypothesized to emerge from stochastic fluctuations in the initial number of cells that take between the two tumors, with variations of about twenty-five percent between the two. Critically, while simulating this difference in initial condition resulted in only small changes when no interaction was taken into account, it generated substantial discrepancies when it was. Our model thus gives a dynamical explanation of the observation of one tumor “winning on the other”, since one and only one of the two tumors gets substantially suppressed despite equal growth and interaction parameters. Notably, within the context of proliferation inhibition, other models such as a cytotoxic (log-kill) effect or production of inhibitory factors by the entire tumor burden had to be rejected. This suggests experimentally testable predictions in the mode of production (only by proliferative cells) and action of the inhibitory factors.

Arguments disregarding the competition theory had already been put forward by others (7,11,40). For example, Gorelik had argued that under this theory, the intensity of CR should be an increasing function of the amount of cells implanted in a subsequent graft, in contradiction with experimental findings, thus disqualifying the theory (11). However, these arguments had remained qualitative. Our formal study adds a quantitative basis to these considerations by showing that, under the modeling assumptions we operated, this theory was unable to accurately describe our data (supplementary Figures 6 and 10A).

More elusive in the literature had remained the question to discriminate between angiogenesis inhibition, as evidenced by the work of Folkman and colleagues (18,32), and direct induction of quiescence by seric factors, as proposed by Ruggiero and colleagues (2,3,13). Our results suggest that the latter theory, when considered alone, could be sufficient to drive CR, insofar as it exhibited good match to the data. However, this does not preclude systemic inhibition of angiogenesis (SIA) to occur, since the two theories are not mutually exclusive. Mutual non-exclusivity also applies to the competition theory: it cannot be completely disregarded that a combination of the three phenomena happens in the occurrence of CR. However, it is beyond the scope of the current study to be able to disentangle between a combination of the phenomena and one phenomenon alone.

In our analysis, we did not address the implication of the immune system in CR, despite the reasonable belief that immune players could be involved. In fact, CR was originally termed “concomitant immunity”, because it was believed that CR occurred principally by triggering an immune response from the presence of the first tumor (7). However, several subsequent studies demonstrated that the phenomenon of CR could occur in a broad range of situations where the immune system could not be the main driver, including studies using the animal model employed here (8,11–13). For example, in (11), the authors demonstrated that, in T-cell-depleted mice bearing Lewis Lung Carcinoma, the CR effect was equally effective as in normal mice, and it was tumor-non-specific. Similar results were obtained when studying CR in mice with functions of macrophages and natural killer cells suppressed (12). Therefore, given the amount of evidence that non-immunological phenomena are involved in the generation of concomitant resistance, and for the sake of simplicity, we decided to focus here on theories that did not require the intervention of immune players and postpone this to future work.

Our findings not only shed light on the dynamics of CR but also proposed a simple and biologically-based model of (double and) single tumor growth able to describe the ubiquitously observed growth retardation with larger volumes (41–44), usually modeled by means of the Gompertz equation (41,45). Although several attempts of deriving the Gompertz equation from basic principles have been performed in the literature (46,47), the model we propose here benefits from its simplicity. It has only two (aggregated) parameters, which quantify two phenomena: 1) proliferation of the active cancer cells and 2) production by these active cells themselves of factors that drive them to quiescence. Interestingly, this model brings new light on the so-called paradox of CR (40), which can be expressed as follows: if distant inhibition occurs, potentially driving other tumors to dormancy, then why does the primary tumor continue growing? Our general model gives a way to quantitatively formalize this. Indeed, the same factors act as local and distant inhibitors and we showed that, under appropriate values of the parameters (but identical for the two tumors), one could obtain at the same time almost unaltered growth of the large (primary) tumor and significant suppression of the growth of the small (secondary) tumor. The presence of endogenous molecules with inhibitory activity thus challenges a naïve view where growth retardation would only be due to interactions dictated by competition (for space or nutrients). Consistently, such a model (logistic growth), had already been shown unable to adequately fit experimental tumor growth curves (36). Considering the implications, the mere fact a tumor would produce both angiogenesis stimulators and inhibitors at the same time, with near and far ranges, does not readily reconcile with a purely localized purpose, but instead speaks to tumor control being manifestly a systemic phenomenon, quite distinct from the naïve concept of an entity governed by local conditions alone, independent of other tumor sites. Implicit in this, and as proposed by others (40,48), a vision of tumor growth as an integrated, organ-like development could bring sense to this seeming paradox.

Concomitant resistance and consequent post-surgery metastatic acceleration (PSMA) have important clinical implications (25–27). In some instances, surgery might not be the optimal therapeutic option and it might be more beneficial to try to control the patient total tumor burden (49). Additionally, it has been experimentally demonstrated that pre-operative administration of chemotherapy or radiotherapy could prevent this acceleration, thus giving a strong rationale for neo-adjuvant therapy (50). Although PSMA has been reported experimentally since more than one century (6), consideration of this phenomenon in the clinic has remained limited. After testing of more than thirty mathematical models of CR, we dispose now of a validated model for the interaction of two tumors. In parallel, we have developed since several years mathematical models for the systemic dynamics of metastases (51–54). Our next step is thus to integrate in these models the non-trivial interactions between the primary tumor and the metastases, or among the metastases themselves, as captured by the mathematical model defined here. When adequately validated, this will provide a computational tool of valuable clinical interest, as it could lead to personalized quantification of the impact of CR in patients. In turn this will bring the opportunity to simulate various therapeutic scenarios and predict the potential occurrence of PSMA, paving the way to patient-specific adaptation of neo-adjuvant and adjuvant therapy.

## Acknowledgments

We thank Dr Sauveur Merlenghi, Dr Marie Dominique Battesti and Mrs N. Spinosi from the Ligue Contre le Cancer de Corse du Sud for their support.

## Supplementary Figures

**Supplementary figure 1:**
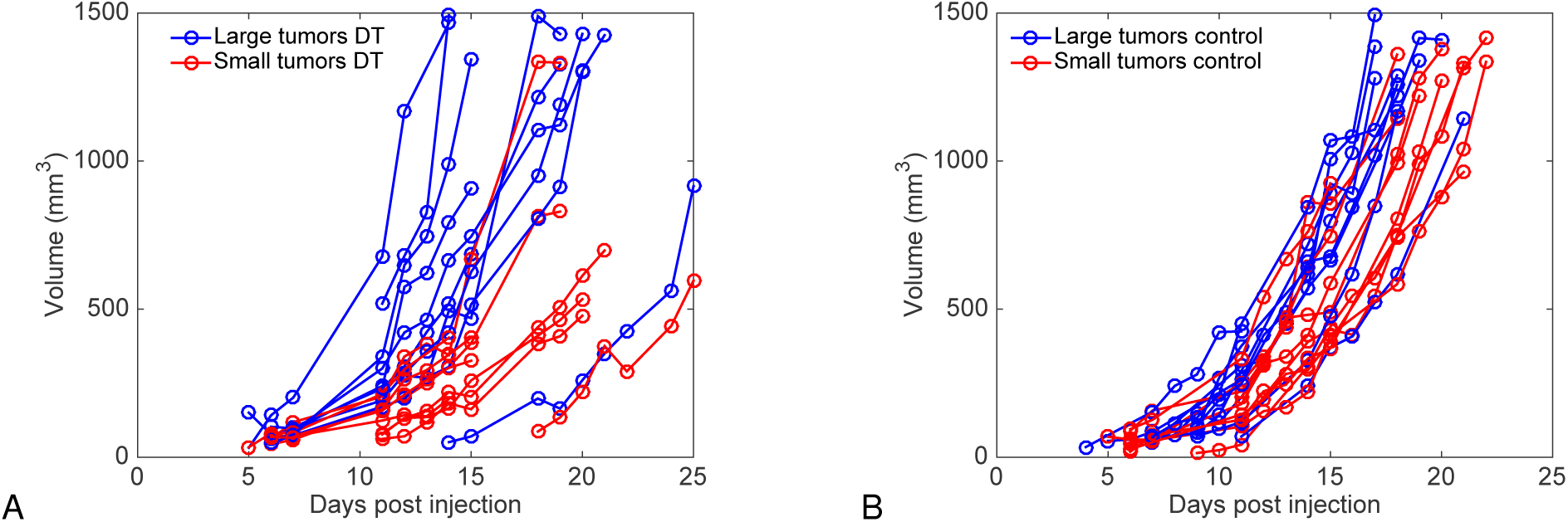
plot of all tumor volumes for double and single (control) tumor groups. Growth curves were plotted and distinguished between large and small tumors for the double (A) and single (control) (B) groups. In the second case, artificial and pairings were considered between tumors of the control group to determine large and small tumors that would have occurred by randomness only (i.e. under independent growth conditions). DT = double tumors group

**Supplementary figure 2:**
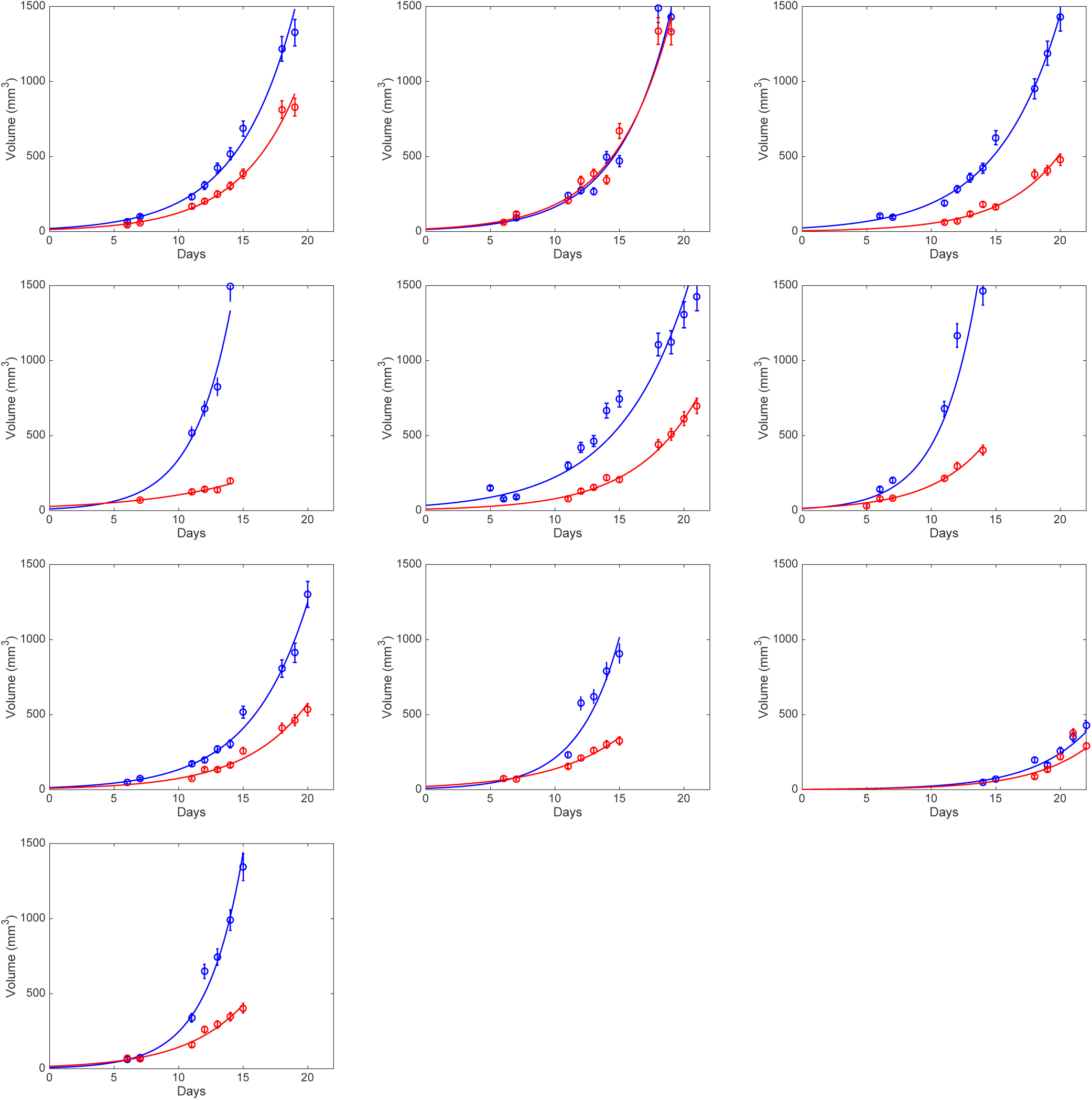
single tumor model fit. Exponential model.

**Supplementary figure 3:**
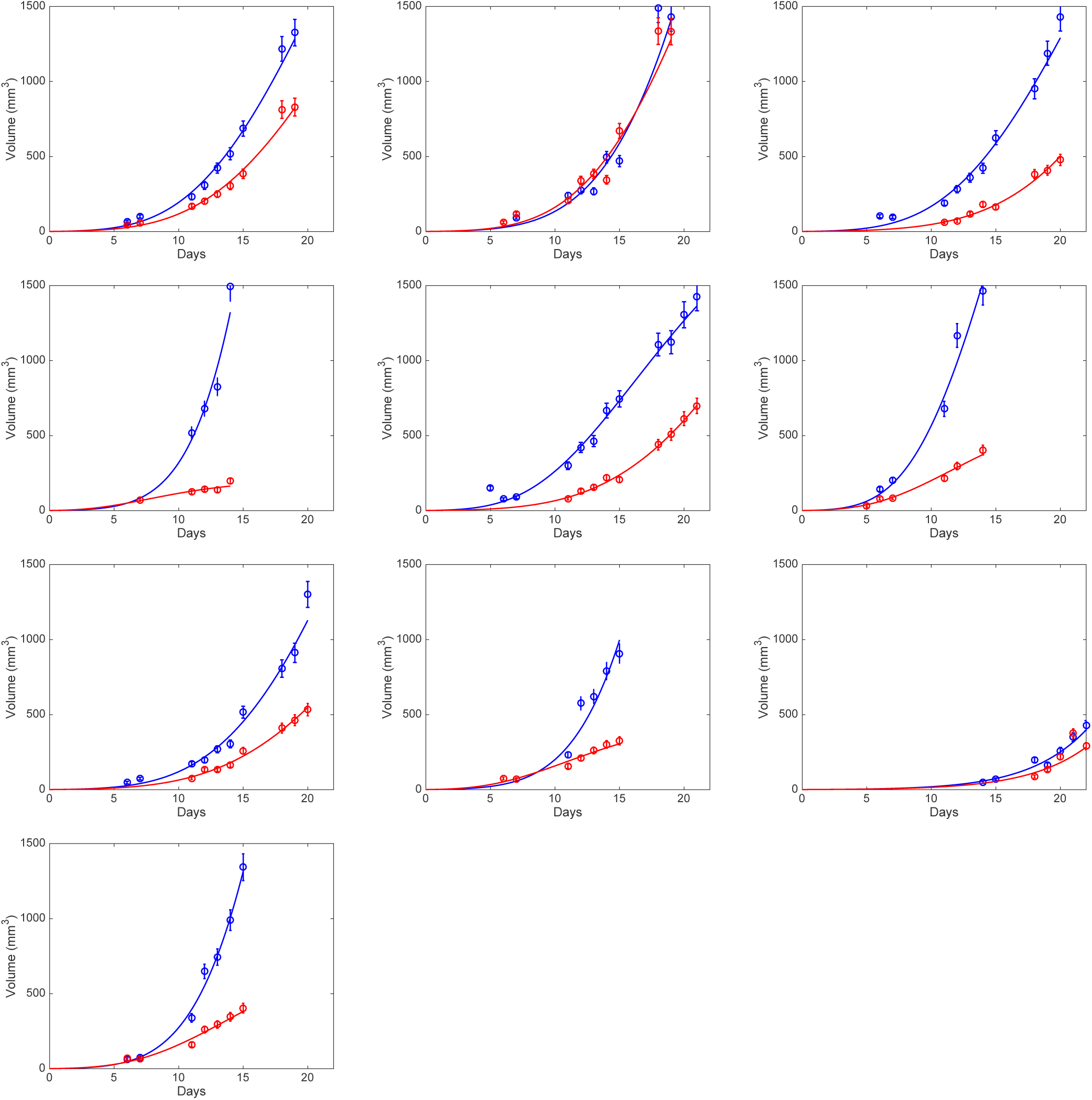
single tumor model fit. Gompertz model.

**Supplementary figure 4:**
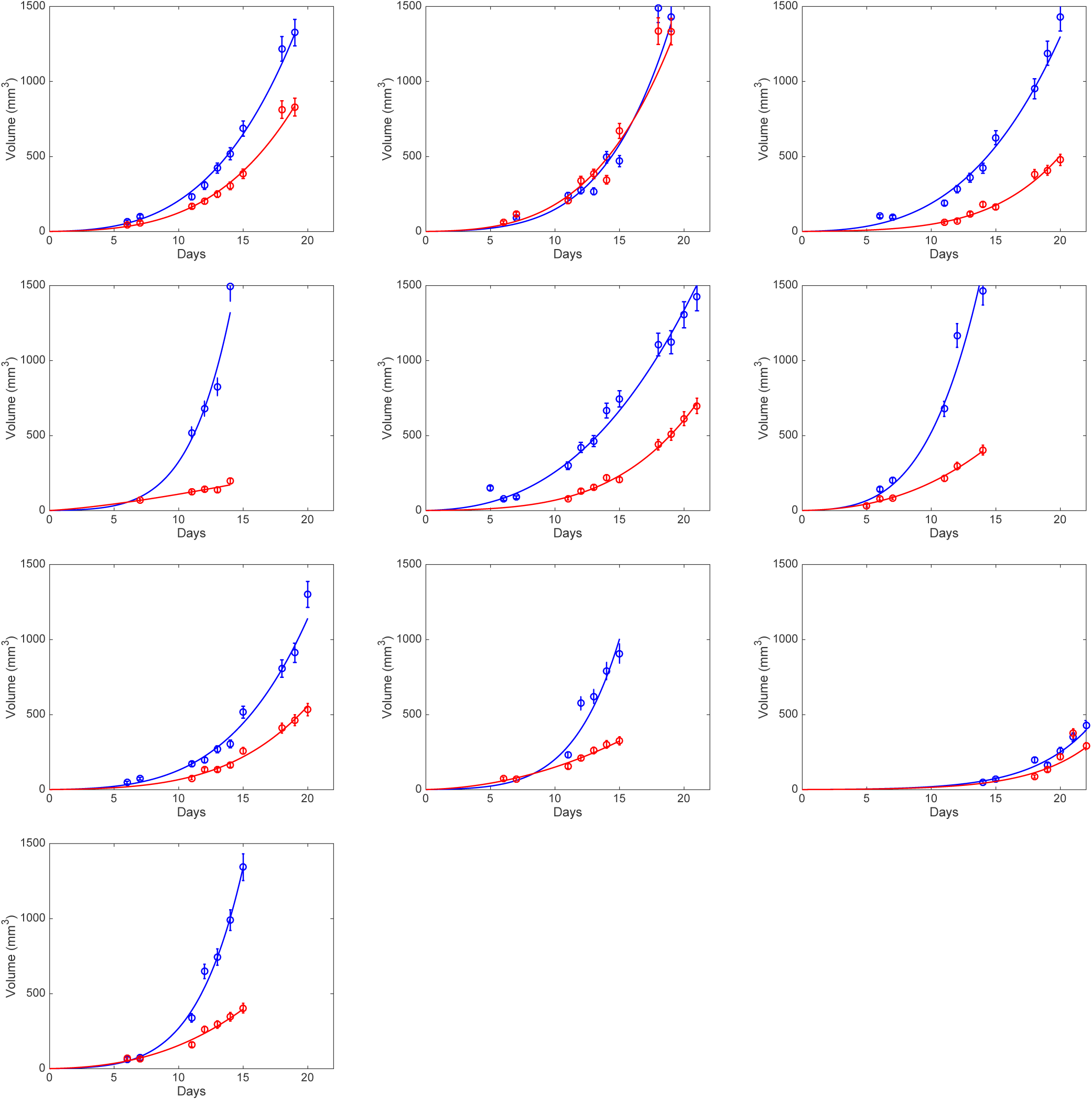
single tumor model fit. Power law model.

**Supplementary figure 5:**
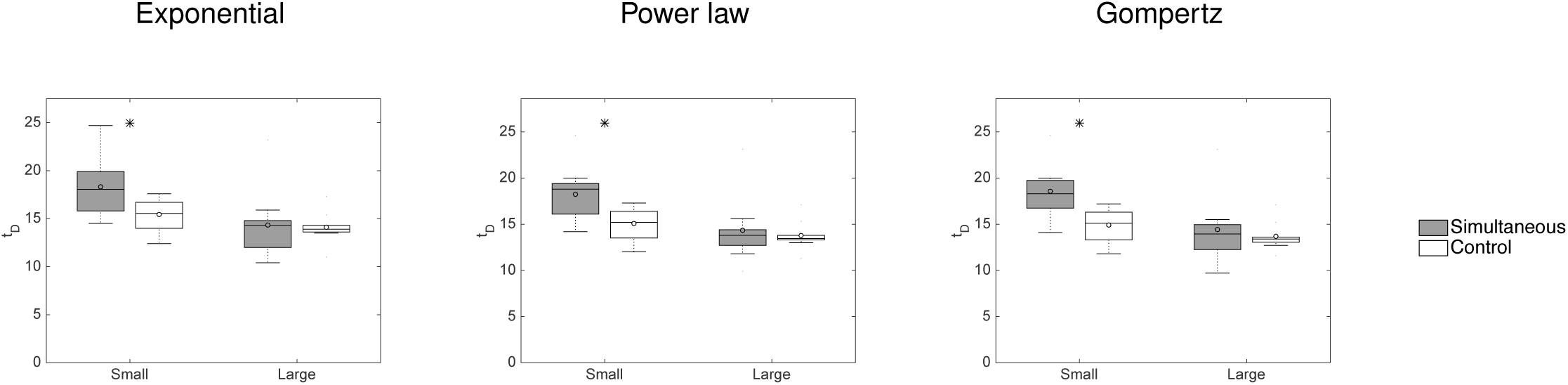
single tumor growth models: delay analysis. Time *t_D_* to reach a threshold volume *V_D_* = 500 mm^3^ for growth extrapolated from the fitted models. * = *p* < 0.05, Student’s t-test with unequal variance.

**Supplementary figure 6:**
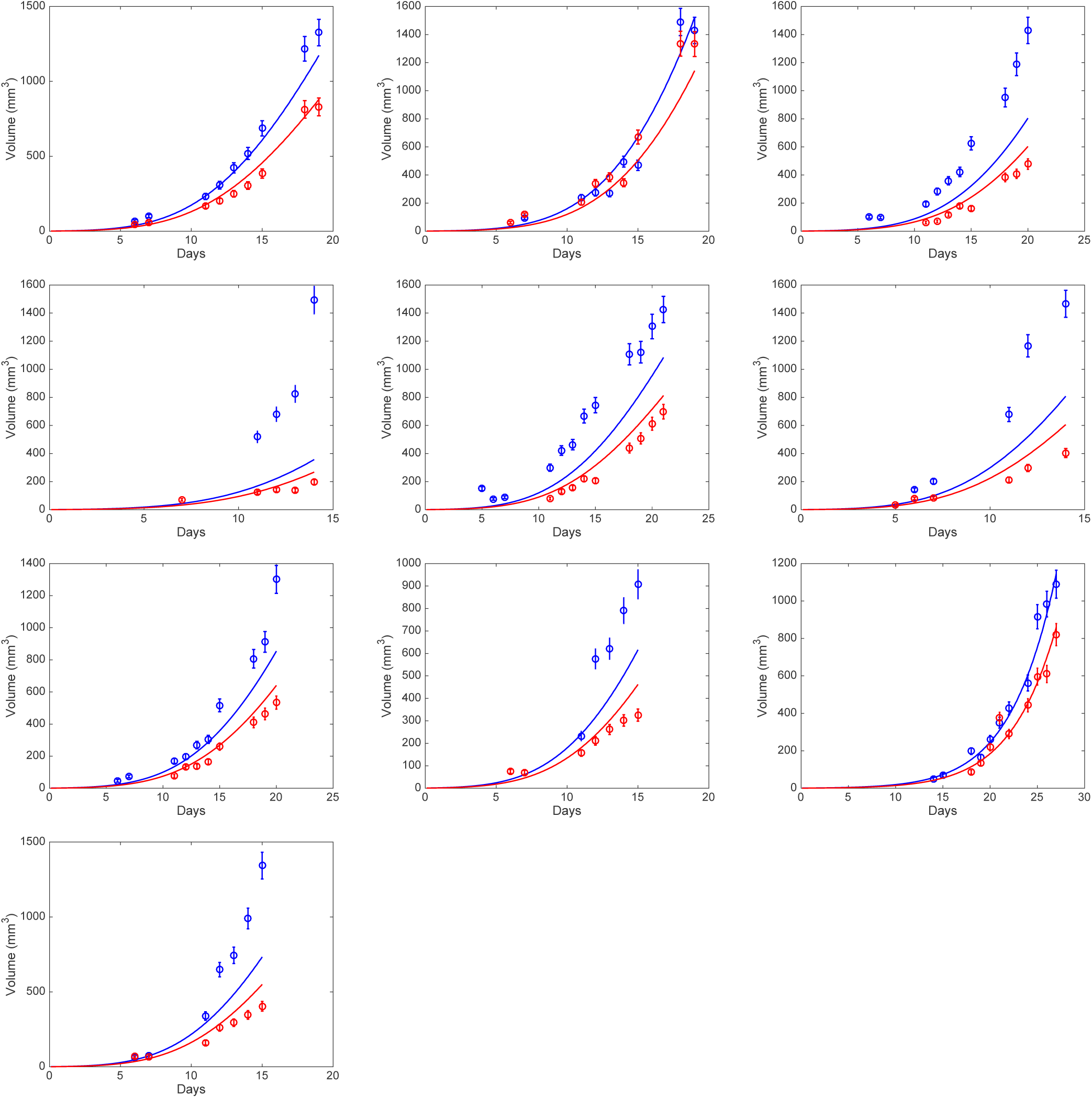
double-tumors growth models fits. Competition model.

**Supplementary figure 7:**
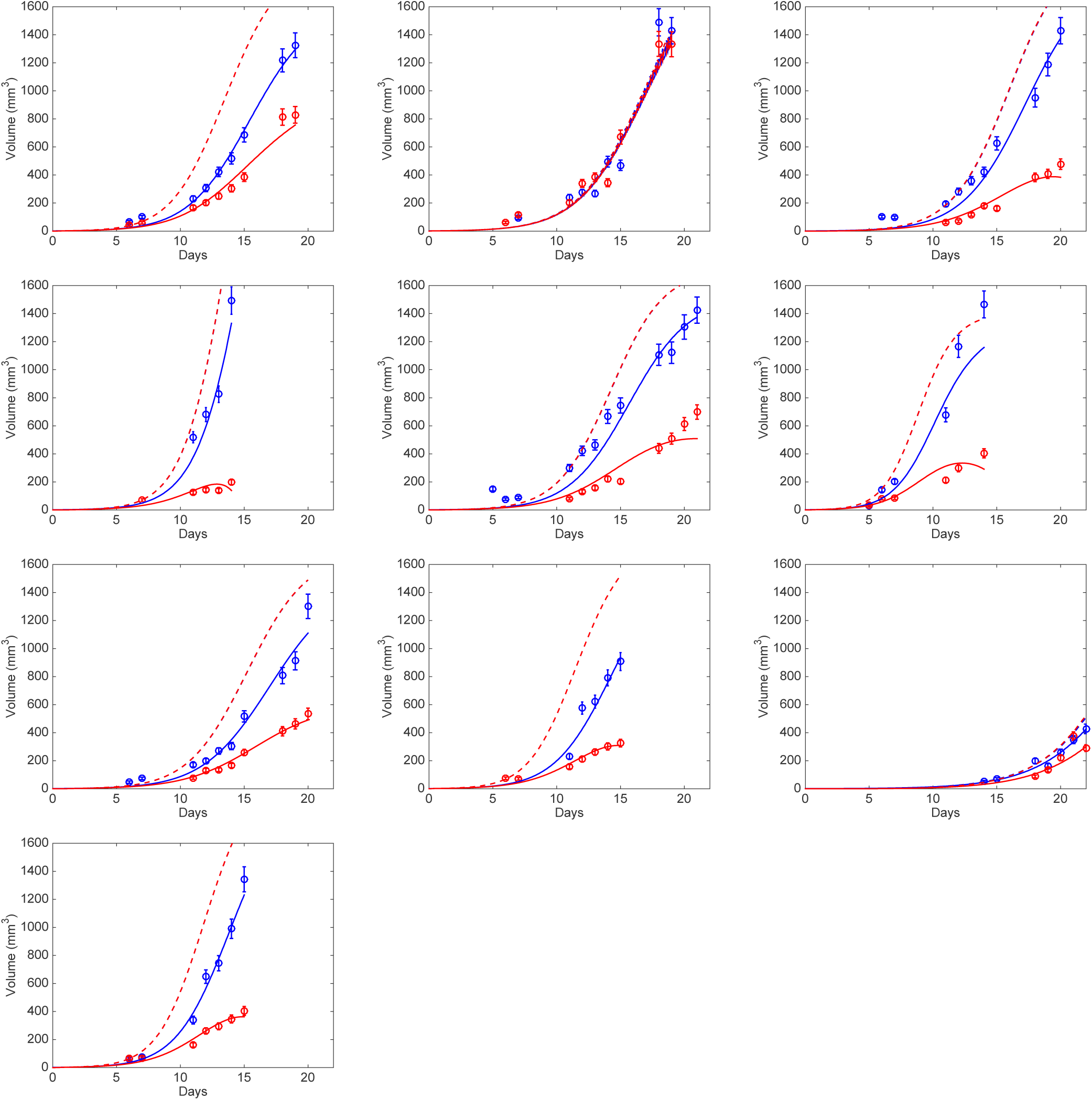
double-tumors growth models fits. Angiogenesis inhibition (SIA) Dashed lines are simulations with no interactions between the two tumors, i.e. with the parameters inferred from the fits except for parameter *e* set to zero. Growth differences are only due to the difference in initial condition.

**Supplementary figure 8:**
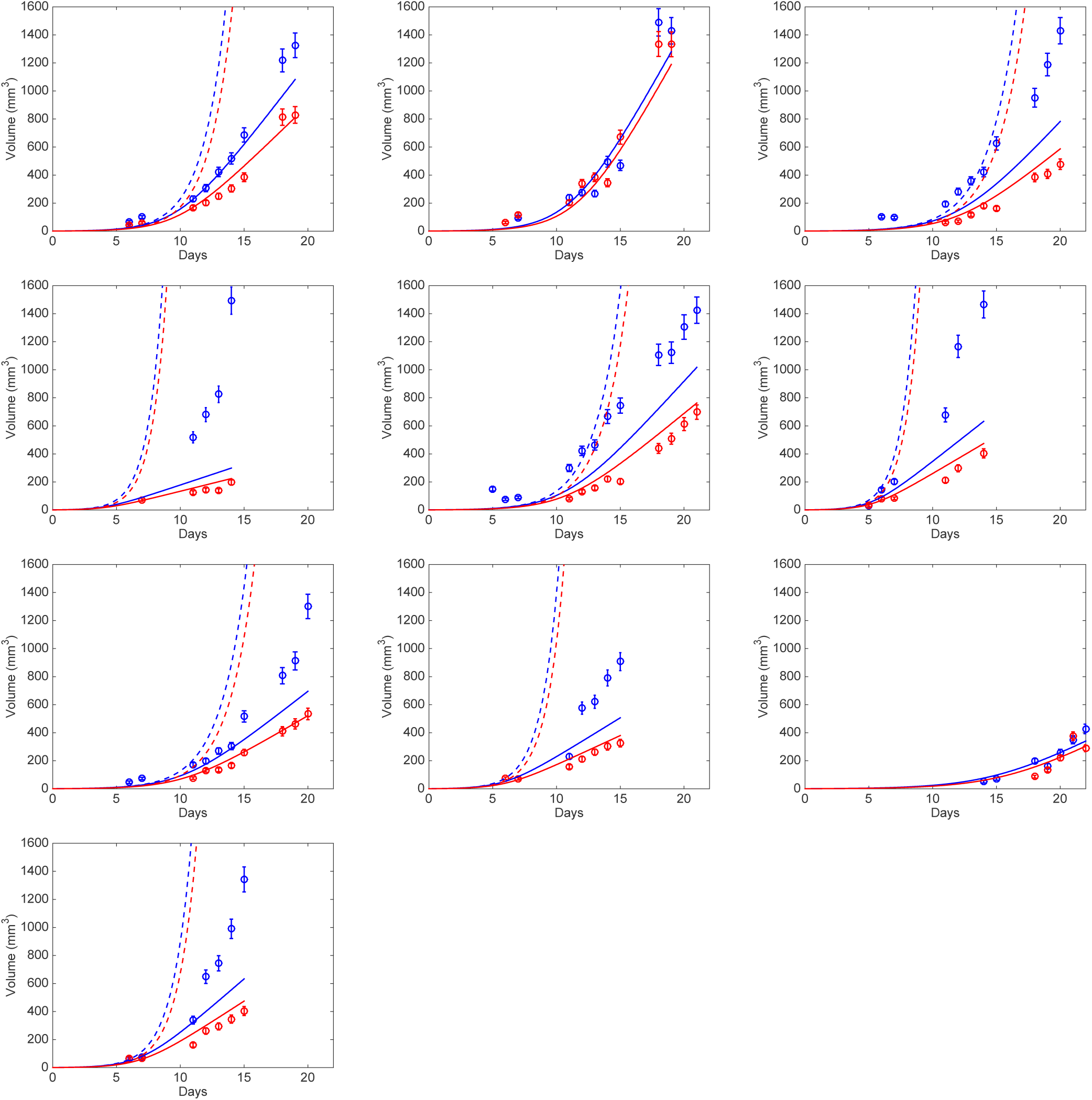
double-tumors growth models fits.Proliferation inhibition. Log-kill effect. Dashed lines are simulations with no interactions between the two tumors. Growth differences are only due to the difference in initial condition.

**Supplementary figure 9:**
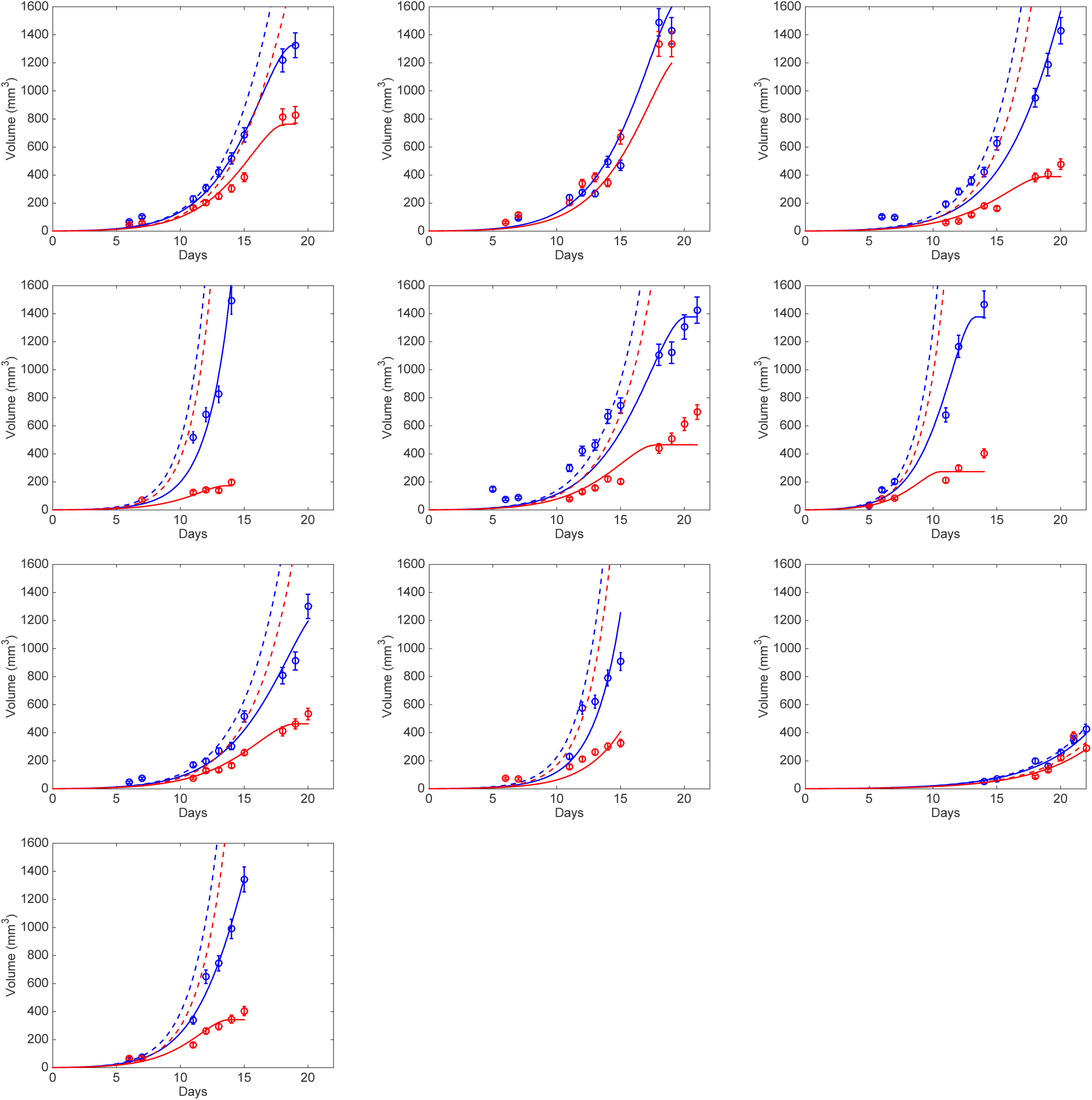
double-tumors growth models fits. Proliferation inhibition. (*P_i_* + *Q_i_*) as source of IFs. Dashed lines are simulations with no interactions between the two tumors. Growth differences are only due to the difference in initial condition.

**Supplementary figure 10:**
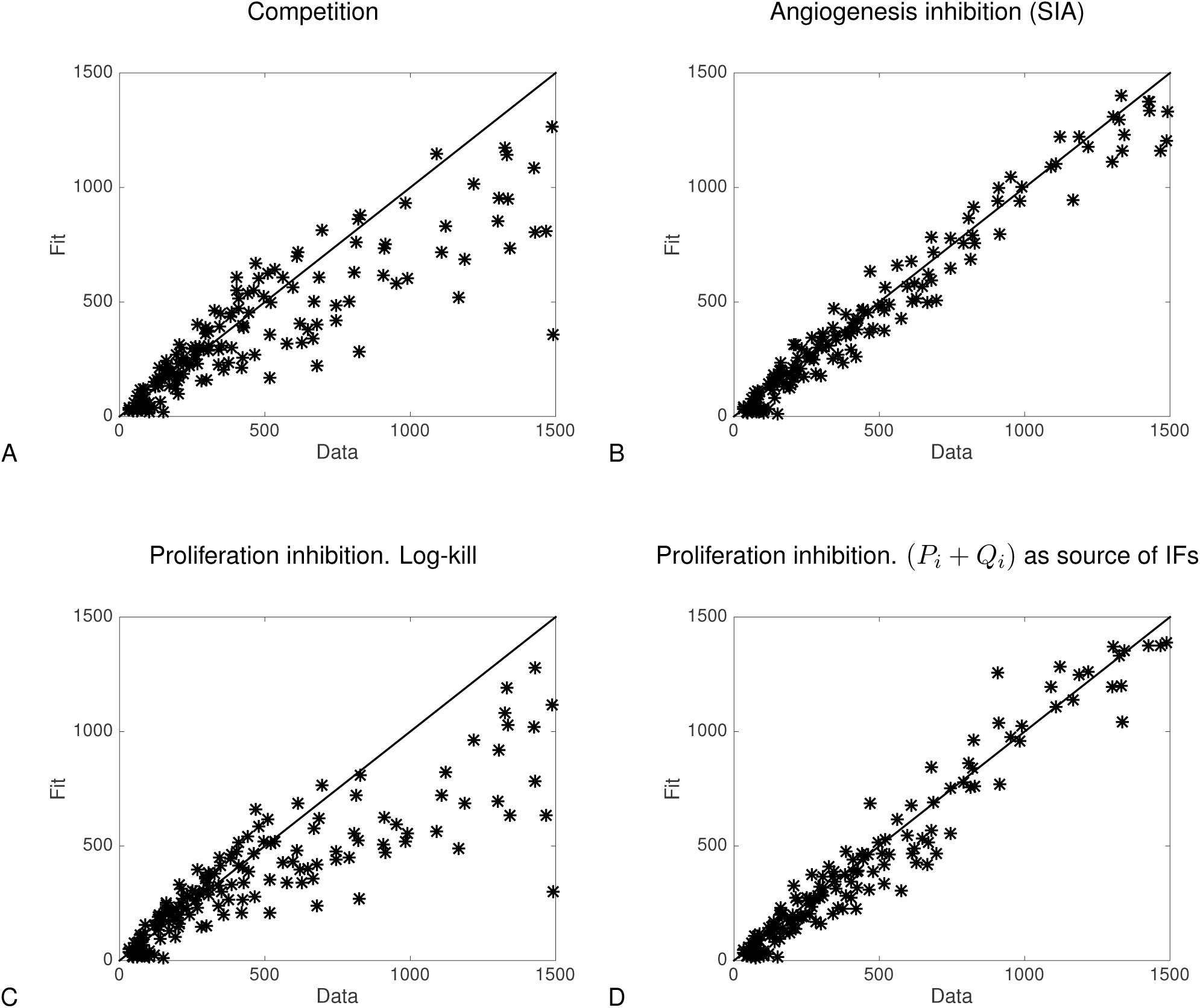
residuals analysis of the other models.

**Supplementary figure 11:**
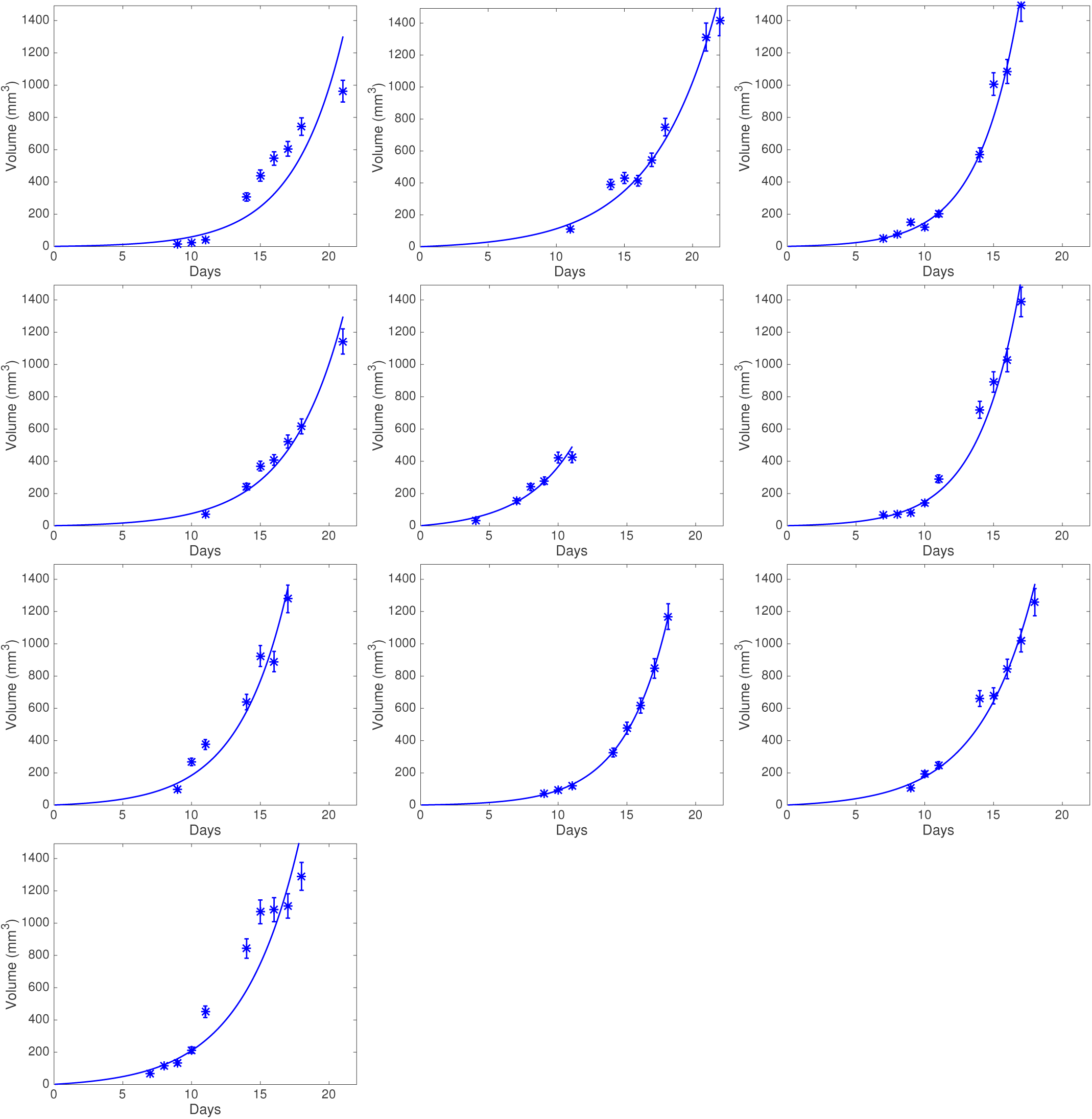
additional fits of 10 remaining animals of the control group under the “proliferation inhibition” model.

**Supplementary table 1:** models equations.

| Model name | Equations |
| --- | --- |
| Competition | $\begin{cases} \frac{dV_1}{dt} = aV_1 \ln\left(\frac{K}{V_1+V_2}\right) & V_1(t=0) = V_{0,1} \\ \frac{dV_2}{dt} = aV_2 \ln\left(\frac{K}{V_1+V_2}\right) & V_2(t=0) = V_{0,2} \end{cases}$ |
| Angiogenesis inhibition | $\begin{cases} \frac{dV_1}{dt} = aV_1 \ln\left(\frac{K_1}{V_1}\right) & V_1(t=0) = V_{0,1} \\ \frac{dK_1}{dt} = bK_1 - dV_1^{2/3}K_1 - eV_2\mathbb{1}_{K_1>K_0} & K_1(t=0) = K_0 \\ \frac{dV_2}{dt} = aV_2 \ln\left(\frac{K_2}{V_2}\right) & V_2(t=0) = V_{0,2} \\ \frac{dK_2}{dt} = bK_2 - dV_2^{2/3}K_2 - eV_1\mathbb{1}_{K_2>K_0} & K_2(t=0) = K_0 \end{cases}$ |
| Proliferation inhibition | $\begin{cases} \frac{dP_1}{dt} = \alpha P_1 - (\beta P_1 + \gamma(P_1 + P_2))\mathbb{1}_{P_1>0} & P_1(t=0) = V_{0,1} \\ \frac{dQ_1}{dt} = (\beta P_1 + \gamma(P_1 + P_2))\mathbb{1}_{P_1>0} & Q_1(t=0) = 0 \\ V_1 = P_1 + Q_1 \\ \frac{dP_2}{dt} = \alpha P_2 - (\beta P_2 + \gamma(P_1 + P_2))\mathbb{1}_{P_2>0} & P_2(t=0) = V_{0,2} \\ \frac{dQ_2}{dt} = (\beta P_2 + \gamma(P_1 + P_2))\mathbb{1}_{P_2>0} & Q_2(t=0) = 0 \\ V_2 = P_2 + Q_2 \end{cases}$ |
| Proliferation inhibition (log-kill) | $\begin{cases} \frac{dP_1}{dt} = \alpha P_1 - (\beta P_1 + \gamma(P_1 + P_2))P_1 & P_1(t=0) = V_{0,1} \\ \frac{dQ_1}{dt} = (\beta P_1 + \gamma(P_1 + P_2)) & Q_1(t=0) = 0 \\ V_1 = P_1 + Q_1 \\ \frac{dP_2}{dt} = \alpha P_2 - (\beta P_2 + \gamma(P_1 + P_2))P_2 & P_2(t=0) = V_{0,2} \\ \frac{dQ_2}{dt} = (\beta P_2 + \gamma(P_1 + P_2)) & Q_2(t=0) = 0 \\ V_2 = P_2 + Q_2 \end{cases}$ |
| Proliferation inhibition ($P+Q$) | $\left\{ \begin{array}{ll} \frac{dP_1}{dt} = \alpha P_1 - (\beta V_1 + \gamma (V_1 + V_2)) \mathbb{1}_{P_1 > 0} & P_1(t=0) = V_{0,1} \\ \frac{dQ_1}{dt} = (\beta V_1 + \gamma (V_1 + V_2)) \mathbb{1}_{P_1 > 0} & Q_1(t=0) = 0 \\ V_1 = P_1 + Q_1 \\ \frac{dP_2}{dt} = \alpha P_2 - (\beta V_2 + \gamma (V_1 + V_2)) \mathbb{1}_{P_2 > 0} & P_2(t=0) = V_{0,2} \\ \frac{dQ_2}{dt} = (\beta V_2 + \gamma (V_1 + V_2)) \mathbb{1}_{P_2 > 0} & Q_2(t=0) = 0 \\ V_2 = P_2 + Q_2 \end{array} \right.$ |

**Supplementary table 2:** goodness-of-fit metrics of these three single tumor growth models.

| Model | SSE | AIC | RMSE | R2 | p |
| --- | --- | --- | --- | --- | --- |
| Power law | **0.117(0.0158 - 0.713)[1]** | **-12.3(-34.5 - 2.95)[1]** | 0.4(0.145 - 0.957)[2] | 0.983(0.784 - 0.998)[2] |  |
| Gompertz | 0.121(0.019 - 0.67)[2] | -11.6(-32.4 - 2.39)[2] | **0.394(0.159 - 0.928)[1]** | **0.984(0.815 - 0.997)[1]** |  |
| Proliferation inhibition | 0.159(0.00741 - 0.883)[3] | -10.1(-33.2 - 4.88)[3] | 0.45(0.0994 - 1.07)[3] | 0.966(0.7 - 0.999)[3] |  |

SSE = Sum of Square Errors, AIC = Akaike Information Criterion, BIC = Bayesian Information Criterion, R2 = coefficient of determination. # = number of parameters.

**Supplementary table 3:** parameter values and identifiability of the Gompertz, power law and “proliferation inhibition” model for the single tumor growth fits.

| Model | Par. | Unit | Median value (CV) | NSE (%) (CV) |
| --- | --- | --- | --- | --- |
| Power law | $\alpha$ | $\text{mm}^{3(1-\gamma)} \cdot \text{day}^{-1}$ | 0.921 (41.9) | 10.6 (55) |
| | $\gamma$ | - | 0.788 (9.35) | 3.42 (62.4) |
| Gompertz | $\alpha_0$ | $\text{day}^{-1}$ | 1.84 (35.7) | 9.28 (65.3) |
| | $\beta$ | $\text{day}^{-1}$ | 0.0792 (43) | 12 (74.4) |
| Proliferation inhibition | $\alpha$ | $\text{day}^{-1}$ | 3.71 (54.4) | 21.4 (61.7) |
| | $\beta + \gamma$ | $\text{day}^{-1}$ | 3.45 (59) | 24.5 (74.5) |

Par. = parameter. CV = coefficient of variation. NSE = normalized standard error.

## References

1. Spratt JS, Meyer JS, Spratt JA. Rates of growth of human solid neoplasms: Part I. J Surg Oncol. 1995;60:137–46.

2. Ruggiero RA, Bruzzo J, Chiarella P, di Gianni P, Isturiz MA, Linskens S, et al. Tyrosine isomers mediate the classical phenomenon of concomitant tumor resistance. Cancer Res. 2011;71:7113–24.

3. Ruggiero RA, Bruzzo J, Chiarella P, Bustuoabad OD, Meiss RP, Pasqualini CD. Concomitant tumor resistance: the role of tyrosine isomers in the mechanisms of metastases control. Cancer Res. 2012;72:1043–50.

4. Skipper HE. The effects of chemotherapy on the kinetics of leukemic cell behavior. Cancer Res. 1965;25:1544–50.

5. Chiarella P, Bruzzo J, Meiss RP, Ruggiero RA. Concomitant tumor resistance. Cancer Lett. 2012;324:133–41.

6. Demicheli R, Retsky MW, Hrushesky WJM, Baum M, Gukas ID. The effects of surgery on tumor growth: a century of investigations. Ann Oncol. 2008;19:1821–8.

7. Gershon RK, Carter RL, Kondo K. On Concomitant Immunity in Tumour-bearing Hamsters. Nature. 1967;213:674–6.

8. Dewys WD. Studies correlating the growth rate of a tumor and its metastases and providing evidence for tumor-related systemic growth-retarding factors. Cancer Res. 1972;32:374–9.

9. Simpson-Herren L, Sanford AH, Holmquist JP. Effects of surgery on the cell kinetics of residual tumor. Cancer Treat Rep. 1976;60:1749–60.

10. Gunduz N, Fisher B, Saffer EA. Effect of surgical removal on the growth and kinetics of residual tumor. Cancer Res. 1979;39:3861–5.

11. Gorelik E, Segal S, Feldman M. On the mechanism of tumor “concomitant immunity”. Int J Cancer. 1981;27:847–56.

12. Gorelik E. Resistance of tumor-bearing mice to a second tumor challenge. Cancer Res. 1983;43:138–45.

13. Ruggiero RA, Bustuoabad OD, Bonfil RD, Meiss RP, Pasqualini, CD. “Concomitant immunity” in murine tumours of non-detectable immunogenicity. Br J Cancer. 1985;51:37–48.

14. Fisher B, Gunduz N, Saffer EA. Influence of the interval between primary tumor removal and chemotherapy on kinetics and growth of metastases. Cancer Res. 1983;43:1488–92.

15. Sckell A, Safabakhsh N, Dellian M, Jain RK. Primary tumor size-dependent inhibition of angiogenesis at a secondary site: an intravital microscopic study in mice. Cancer Res. 1998;58:5866–9.

16. Li TS, Kaneda Y, Ueda K, Hamano K, Zempo N, Esato K. The influence of tumour resection on angiostatin levels and tumour growth--an experimental study in tumour-bearing mice. Eur J Cancer. 2001;37:2283–8.

17. Marie P, Clunet J. Frequency of visceral metastases in tumor bearing mice following surgical removal of their tumor [French]. Bull Assoc Fr Etud Cancer. 1910;3:19–23.

18. O’Reilly MS, Holmgren L, Shing Y, Chen C, Rosenthal RA, Moses M, et al. Angiostatin: a novel angiogenesis inhibitor that mediates the suppression of metastases by a Lewis lung carcinoma. Cell. 1994;79:315–28.

19. Rofstad E, Graff B. Thrombospondin-1-mediated metastasis suppression by the primary tumor in human melanoma xenografts. J Invest Dermatol. 2001;117:1042–9.

20. Gorelik E, Segal S, Feldman M. Growth of a local tumor exerts a specific inhibitory effect on progression of lung metastases. Int J Cancer. 1978;21:617–25.

21. Peeters CF, de Waal RM, Wobbes T, Ruers TJ. Metastatic dormancy imposed by the primary tumor: does it exist in humans? Ann Surg Oncol. 2008;15:3308–15.

22. Ketcham AS, Kinsey DL, Wexler H, Mantel N. The development of spontaneous metastases after the removal of a “primary” tumor. II. Standardization protocol of 5 animal tumors. Cancer. 1961;14:875–82.

23. Fisher B, Gunduz N, Coyle J, Rudock C, Saffer E. Presence of a growth-stimulating factor in serum following primary tumor removal in mice. Cancer Res. 1989;49:1996–2001.

24. Ceelen W, Pattyn P, Mareel M. Surgery, wound healing, and metastasis: recent insights and clinical implications. Crit Rev Oncol Hematol. 2014;89:16–26.

25. Peeters CFJM, de Waal RMW, Wobbes T, Westphal JR, Ruers TJM. Outgrowth of human liver metastases after resection of the primary colorectal tumor: a shift in the balance between apoptosis and proliferation. Int J Cancer. 2006;119:1249–53.

26. Demicheli R, Retsky MW, Hrushesky WJM, Baum M. Tumor dormancy and surgery-driven interruption of dormancy in breast cancer: learning from failures. Nat Clin Rev Oncol. 2007;4:699–710.

27. Coffey JC, Wang JH, Smith MJF, Bouchier-Hayes D, Cotter TG, Redmond HP. Excisional surgery for cancer cure: therapy at a cost. Lancet Oncol. 2003;4:760–8.

28. Retsky M, Demicheli R. Multimodal Hazard Rate for Relapse in Breast Cancer: Quality of Data and Calibration of Computer Simulation. Cancers. 2014;6:2343–55.

29. Demicheli R, Abbattista A, Miceli R, Valagussa P, Bonadonna G. Time distribution of the recurrence risk for breast cancer patients undergoing mastectomy: Further support about the concept of tumor dormancy. Breast Cancer Res Treat. 1996;41:177–85.

30. Ehrlich P. Experimentelle Carcinomstudien an Mäusen. Arb Aus Dem K Inst Für Exp Ther Zu Frankf Am Main. 1906;75–102.

31. O’Reilly MS, Boehm T, Shing Y, Fukai N, Vasios G, Lane WS, et al. Endostatin: an endogenous inhibitor of angiogenesis and tumor growth. Cell. 1997;88:277–85.

32. Holmgren L, O’Reilly MS, Folkman J. Dormancy of micrometastases: balanced proliferation and apoptosis in the presence of angiogenesis suppression. Nat Med. 1995;1:149–53.

33. Folkman J. Angiogenesis in cancer, vascular, rheumatoid and other disease. Nat Med. 1995;1:27–31.

34. Bertram JS, Janik P. Establishment of a cloned line of Lewis Lung Carcinoma cells adapted to cell culture. Cancer Lett. 1980;11:63–73.

35. Kunstyr I, Leuenberger HG. Gerontological data of C57BL/6J mice. I. Sex differences in survival curves. J Gerontol. 1975;30:157–62.

36. Benzekry S, Lamont C, Beheshti A, Tracz A, Ebos JML, Hlatky L, et al. Classical mathematical models for description and prediction of experimental tumor growth. Mac Gabhann F, editor. PLoS Comput Biol. 2014;10:e1003800.

37. Seber GA, Wild CJ. Nonlinear regression. Wiley-Interscience; 2003.

38. d’Onofrio A, Gandolfi A. Tumour eradication by antiangiogenic therapy: analysis and extensions of the model by Hahnfeldt et al. (1999). Math Biosci. 2004;191:159–84.

39. Gompertz B. On the Nature of the Function Expressive of the Law of Human Mortality, and on a New Mode of Determining the Value of Life Contingencies. Phil Trans R Soc B. 1825;115:513–83.

40. Prehn RT. Two competing influences that may explain concomitant tumor resistance. Cancer Res. 1993;53:3266–9.

41. Laird AK. Dynamics of tumor growth. Br J Cancer. 1964;13:490–502.

42. Steel GG, Lamerton LF. The growth rate of human tumours. Br J Cancer. 1966;20:74–86.

43. Spratt JA, von Fournier D, Spratt JS, Weber EE. Decelerating growth and human breast cancer. Cancer. 1993;71:2013–9.

44. Wheldon TE. Mathematical models in cancer research. Hilger Bristol; 1988.

45. Norton L. A Gompertzian model of human breast cancer growth. Cancer Res. 1988;48:7067–71.

46. Frenzen CL, Murray JD. A Cell Kinetics Justification for Gompertz’ Equation. SIAM J Appl Math. 1986;46:614–29.

47. Bajzer Ž, Vuk-Pavlović, S, Huzak M. Mathematical modeling of tumor growth kinetics. Surv Models Tumor-Immune Syst Dyn. Springer; 1997. page 89–133.

48. Prehn RT. The inhibition of tumor growth by tumor mass. Cancer Res. 1991;51:2–4.

49. Badwe R, Parmar V, Hawaldar R, Nair N, Kaushik R, Siddique S, et al. Surgical removal of primary tumor and axillary lymph nodes in women with metastatic breast cancer at first presentation: A randomized controlled trial. San Antonio Breast Cancer Symp. 2013. page S2–2.

50. Fisher B, Saffer E, Rudock C, Coyle J, Gunduz N. Effect of Local or Systemic Treatment Prior to Primary Tumor Removal on the Production and Response to a Serum Growth-stimulating Factor in Mice. Cancer Res. 1989;49:2002–4.

51. Benzekry S, André N, Benabdallah A, Ciccolini J, Faivre C, Hubert F, et al. Modelling the impact of anticancer agents on metastatic spreading. Math Model Nat Phenom. 2012;7:306–36.

52. Benzekry S, Gandolfi A, Hahnfeldt P. Global Dormancy of Metastases Due to Systemic Inhibition of Angiogenesis. PLoS ONE. 2014;9:e84249–11.

53. Benzekry S, Tracz A, Mastri M, Corbelli R, Barbolosi D, Ebos JML. Modeling spontaneous metastasis following surgery: an in vivo-in silico approach. Cancer Res. 2015;canres.1389.2015.

54. Baratchart E, Benzekry S, Bikfalvi A, Colin T, Cooley LS, Pineau R, et al. Computational Modelling of Metastasis Development in Renal Cell Carcinoma. PLoS Comput Biol. 2015;11:e1004626.

